# Endoderm Nitric Oxide Signals to Regulate Nascent Development of Cardiac Progenitors in Chicken Embryos

**DOI:** 10.1101/2020.08.29.272989

**Authors:** Devan H. Shah, Sujoy K. Biswas, Adrian M. Martin, Simone Bianco, Wilfred F. Denetclaw

## Abstract

Heart development in the chicken embryo is regulated by a concert of cardiogenic morphogens and signaling molecules, but the physiological signal molecule nitric oxide(NO) has not been studied in the context of heart formation. A dynamic investigation of endoderm NO formation demonstrates for the first time a correlation with the established development events of the cardiac heart fields and heart tube. Manipulation of endoderm NO signaling demonstrate a role of NO signaling in the differentiation and proliferation of cardiac progenitors for heart tube formation and cardiac heart field development. To investigate NO in the proliferation of myocardial cells in the heart tube embryos, a computer vision based artificial intelligence approach is followed to automate the long and tedious job of counting cells in a large image dataset. We document NO as an important signaling molecule in the regulation of nascent embryonic cardiogenesis whose effects on other early cardiogenic morphogens is unknown.

## INTRODUCTION

The first organ to develop in amniote embryos is the heart, a contractile tube that soon integrates with vascular tissue making the primitive system (Stalsberg and DeHaan, 1969; Dyer and Kirby, 2010). The inception of cardiogenesis occurs before Hensen’s Node regression in gastrulation as mesoderm cardiac precursor cells ingress and migrate to either lateral side from the primitive streak forming the pre-cardiac mesoderm between endoderm and ectoderm germ layers, called the bilateral heart fields (HF) (Rosenquist, 1970; Garcia-Martinez and Schoenwolf, 1993; Psychoyos and Stern, 1996). The HFs contain cardiac precursor cells specified and determined to cardiac lineage fates as the closely associated endoderm and splanchnic mesoderm coalesce along the embryonic midline to fold upon themselves and to signal with cardiogenic inductive signals.(Yatskievych, Ladd and Antin, 1997; Ladd, Yatskievych and Antin, 1998; Lopez-Sanchez *et al*., 2009) The folded splanchnic mesoderm forms bilateral epithelia comprised of a heterogenous population of cardiac progenitor/precursor cells which give rise to two cell types, endocardial epithelial cells and cardiomyocytes. (Cohen-Gould and Mikawa, 1996; Bussmann, Bakkers and Schulte-Merker, 2007) The first HF in the bilateral HFs develops into endocardial tubes, consisting of a thin outer myocardium and inner tube of endocardial epithelium tube separated by acellular cardiac jelly. (Virágh, Szabó and Challice, 1989; Ivanovitch, Esteban and Torres, 2017). Further Subsequent heart development produces a singular heart tube by the merging and fusion of the paired bilateral endocardial tubes and a complete surrounding of cardiomyocytes at the embryo midline facilitated by an posterior elongating anterior intestinal portal in forming the foregut. (Stalsberg and DeHaan, 1969; Wittig and Münsterberg, 2016). Subsequent elongation of the nascent heart tube that is the left ventricle and partial right ventricle consists of addition of second HF cardiogenic cells to the anterior and posterior poles of the tubular heart to complete the right ventricle and outflow tract and the left and right atrial chambers and inflow tract, respectively. (Kelly, Buckingham and Moorman, 2014; Hosseini, Garcia and Taber, 2017; Kidokoro *et al*., 2018) Shortly after a heart looping event occurs to pre-structure the four chambered heart in the adult animal. (Taber *et al*., 2014; Laura *et al*., 2020)

Nitric Oxide (NO), once considered to be an environmental pollutant and toxic to humans, now is recognized as a gaseous free radical, messenger molecule with many functions in both normal physiology and pathophysiology states. (Ignarro, 2019) NO was first recognized in a biological context to regulate by a soluble guanylyl cyclase (sGC) and cGMP pathway blood vessel vascular tone, local blood flow, and blood pressure (Arnold *et al*., 1977; Furchgott and Zawadzki, 1980; Ignarro *et al*., 1980; Furchgott, 1996). NO research has shown pleiotropic NO signaling functional in many organ systems from the cardiovascular to the muscle, and neural systems and mediated by its lineage specific cell-types (Hibbs et al., 1988; Bredt and Snyder, 1994; Napoli et al., 2013; Tomankova, Abaffy and Sindelka, 2017). NO signals through a sGC/cGMP canonical pathway where the cGMP messenger molecules regulates downstream effectors like Protein Kinase G(PKG). (Quinn, Petros and Vallance, 1995; Cary, Winger and Marletta, 2005). NO is required for normal embryo development (Gouge *et al*., 1998),and has regulatory functions in (Jacox *et al*., 2014), epidermis formation (Tomankova, Abaffy and Sindelka, 2017), neural tube closure (Nachmany *et al*., 2006), skeletal myogenesis (Cazzato *et al*., 2014) and in other developmental events in early embryogenesis. In amniotes, three conserved nitric oxide synthases (NOS1, or neural NOS; NOS2, or inducible NOS; NOS3, or endothelial NOS) produce NO at different rates and durations and under different conditions. (Martínez-Ruiz, Cadenas and Lamas, 2011). However, NO signaling in embryo cardiogenesis and heart development has not been investigated in higher vertebrate embryos although *in vitro* studies of mouse and human ESCs suggest a NO signaling role in cardiac lineage specification and determination of cells(Kim *et al*., 2004; Nikonoff *et al*., 2008).

Cardiogenic specification and determination in the bilateral heart fields depends on the molecular signaling from secreted BMPs, Wnts, and Shh morphogens and their binding proteins, and growth factors, like FGF8 and HGF (Rappolee, Iyer and Patel, 1996; Schultheiss, Burch and Lassar, 1997; Chapman *et al*., 2002; Jiao *et al*., 2006; Goddeeris *et al*., 2007; Klaus *et al*., 2007). These signaling molecules are regulated to produce in the HFs a complex spatial-temporal gradient that defines embryonic regions of lateral cardiac promoting signals and medial regions of cardiac inhibitory signals. (Brade *et al*., 2013; Kelly, Buckingham and Moorman, 2014) Cardiac precursor cells then differentiate in appropriate HF locations to cardiogenic progenitors and subsequent cardiomyocytes and endocardial cells (Alexandra Klaus *et al*., 2007; Münsterberg and Yue, 2008; Tirosh-Finkel *et al*., 2010; Klaus *et al*., 2012). Lateral endoderm derived cardiogenic inductive morphogens like FGF8, which through *Erk* activates MAPK signaling for expression of target genes *Nkx-2.5* and *Mef2c*, (Alsan and Schultheiss, 2002; Jiao *et al*., 2006) and BMP2/4, which activates its effectors smad4 and smad1/5/8 to induce expression of downstream target genes *Gata4* and *Isl1* (Ladd, Yatskievych and Antin, 1998; A. Klaus *et al*., 2007), signal to differentiate cardiac progenitors in the underlying lateral splanchnic mesoderm within the HF. These morphogens signal for cardiac specification and differentiation by inducing expression of transcription factors like *Nkx-2.5/Gata4/Isl1/Mef2c* which act as central regulators of the nascent cardiogenic program (Alsan and Schultheiss, 2002; Chapman *et al*., 2002; Jiao *et al*., 2006; Lopez-Sanchez *et al*., 2009; Koenig *et al*., 2015). Conversely, at the embryo midline, morphogens like the canonical Wnts 1&3, which activate nuclear translocation of β-catenin for transcription of genes involved in cellular proliferation like *c-myc* and *Bcl2* (Alexandra Klaus *et al*., 2007; Tzahor, 2007), and Shh, which induces nuclear translocation of effector GLiR for transcription of target genes involved in proliferation like *Patched* and *Smoothened* (López and Carrasco, 2008; Dyer and Kirby, 2009), signal to maintain second heart field progenitors in an undifferentiated state and restrict pro-cardiogenic signaling to the lateral first HF regions. In the bilateral HFs medial regions, defined by molecular signaling events to delay cardiac differentiation and signal for cellular proliferation, are called the secondary HF (Marvin *et al*., 2001a; Münsterberg and Yue, 2008; Klaus *et al*., 2012). Though advances in molecular signaling have given a model of cardiogenesis in development of the first and secondary HFs, little information is known about NO participation in nascent cardiac development.

Investigations of NO signaling in cardiogenesis has been investigated in multiple *in vitro* systems. Mouse and human ES cells have been directed to cardiogenic lineage fates through the NO canonical signaling pathway to suggest that NO could regulate early embryo cardiogenesis (Kim *et al*., 2004; Mujoo, Krumenacker and Wada, 2006; Madhusoodanan and Murad, 2007; Beyer *et al*., 2015) Key cardiac transcription factors *Nkx-2.5, Gata4,* and *Tbx* mRNA expression upregulate under NO canonical signaling through sGC/cGMP pathway (Kim *et al*., 2004; Mujoo *et al*., 2008; Tejedo *et al*., 2010) Furthermore, eNOS mediated NO production in neonatal mouse cardiac progenitors cells supports proliferation and produces high levels of cardiomyocyte differentiation *in vitro* embryoid body culture and *in vivo* postnatal mouse development. (Lepic *et al*., 2006; Saura *et al*., 2011; Friart *et al*., 2016) Potent effects of NO on embryonic cardiomyogenesis is also shown by differentiation arrest by decreased expression of cardiac muscle proteins, like a-actinin and troponin-1, in response to NOS inhibition in mouse and rat embryoid bodies as well as in the development of the multichambered heart *in vivo*. (Bloch *et al*., 1999; Kim *et al*., 2004; Nikonoff *et al*., 2008; Liu and Feng, 2012). However, NO signaling during early embryo development is poorly studied and has not been directly investigated during the stages of cardiogenesis leading to the formation of the embryo heart tube. Furthermore, *in vitro* investigations of NO regulation of early cardiac lineage development do not give insight to NO spatial and temporal signaling involved in early stages of heart formation. The chicken (*Gallus Gallus domesticus*) embryo has been demonstrated as an ideal model for studying of cardiogenesis at the earliest times of its development because the chicken embryo may develop ex ovo with the embryo dorsally located above the yolk, easily accessible for experimental manipulation and live imaging. Here we use the chicken embryo model to investigate NO signaling patterns during nascent embryonic development and its effects on the differentiation and proliferation of cardiac progenitors in the development of the HFs and formation of the heart. We show for the first time, a pattern of NO signaling in the cardiogenic signaling endoderm correlating to the first and secondary HF, as well as a necessary role of endoderm NO signaling for *in vivo* development of cardiac progenitor cells in higher vertebrate heart tube formation.

## RESULTS

### Early embryo cardiogenesis is signaled by Endoderm NO

We show here chicken embryos from development stages HH3 to HH11 monitored for NO using the irreversible fluorescent DAF2 NO probe in endoderm and ectoderm germ layers adjacent to cardiac mesoderm undergoing development into the bilateral heart fields where the differentiation of cardiogenic progenitor cells into cardiomyocytes takes place and subsequently migrates to the embryo midline to fuse beginning the formation of the heart tube. To investigate NO elevation in epithelial cell layers in areas of cardiogenic mesoderm of the bilateral heart fields, embryos were imaged in both ventral endoderm and dorsal ectoderm layers (Fig1 A-C). When comparing the DAF2 fluorescence intensity of endoderm and ectoderm, normalizing to the average fluorescence, the ectoderm fluorescence was almost 2.5-fold lower than endoderm and the difference is significant (Fig1E). The greater prevalence in endoderm NO signaling to cardiac mesoderm was then monitored for NO changes during stages of early heart formation. Our findings show NO signaling patterns coordinated with nascent development of bilateral heart fields and heart tube formation.

**Figure 1.**
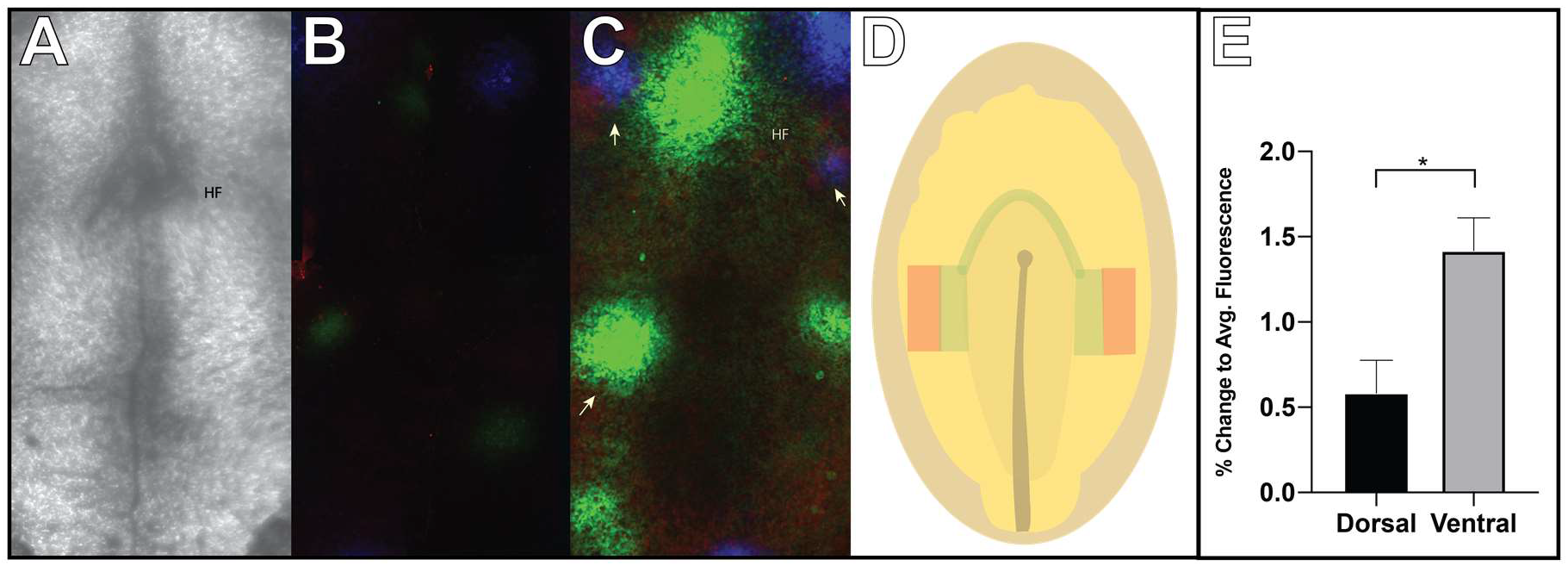
NO is enriched in ventral endoderm. (A) Ventral transmitted light image of a HH5-HH6 embryo to show endoderm over the forming heart field (HF). Embryos prepared as described in Materials and Methods 1.2 were imaged ventrally, and dorsally for DAF2-DA fluorescence (maximum Z-projections of 10 stacks with 10 uM spacing). Composite DAF2-DA fluorescence images of three embryos of comparable ages (red, blue, green representative images of separate embryos) at (B) the dorsal ectoderm which produced NO to a lesser degree (0.583±0.194 A.U. n=3) compared to (C) ventral endoderm (1.417±0.194 A.U. n=3)., which produced NO hotspots at high intensity (arrows) (D) Cartoon of HH5-6 embryo with first heart field progenitors (green), and second heart field progenitors (red) which may correlate to the NO hotspots. (E) Semi-quantitative analysis was performed to compare DAF2-DA fluorescence signal demonstrating significantly less dorsal ectoderm NO signaling compared to ventral endoderm, normalized to a mean average fluorescence between ventral and dorsal images per embryo. One-tailed Student’s t-test. (*p<0.02).

To determine more closely the spatial and temporal signaling of NO in correlation with the development of the HFRs, embryos at specific development stages were monitored by DAF2 fluorescence. In the nascent heart fields of HH4-5 (n=12) staged embryos, the endoderm exhibited a variety of signaling patterns. We observed focal spots of NO signaling in the anterior bilateral heart fields, as shown in orange, white, and green. (Figure 2 A-C). However, we also observed diffuse NO signaling across the lateral endoderm, as shown in magenta, yellow, and blue without evident polarity, suggesting a dynamic activity in NO signaling (Figure 2 C & Supp. Figure 1). In HH5-6 (Fig1 C & Supp. Figure 1) staged embryos (n=5), we observed two patterns of NO signaling where there was a noticeable large diffuse pattern of low level NO in the endoderm overlying the HF (red) and also places of highly elevated and narrowed NO signal we describe as focal hotspots (green and blue) that overly the anterior bilateral HFRs where heart formation is beginning. NO formation was observed to become more restricted to medial and lateral aspects of the endoderm overlying the HFRs, coincident with the overlying differentiating cardiac mesoderm. Endocardial tube (ET) formation (HH7), the folding HFRs and descending anterior intestinal portal tissues were highly elevated in NO (Figure 2 E-G & & Supp. Figure 3). **(n=12)** In these embryos, we also observed broad elevation of NO in the endoderm overlying the splanchnic mesoderm where cardiac progenitor cells and cardiomyocytes were migrating towards the embryo mid-line in formation of the endocardial tubes(green, cyan, red, and yellow). For endocardial tube fusion and primitive heart tube formation **(HH9-10)**, NO formation is abundant in the epicardium overlying the head and fusing ETs, (Figure 2 I-K)**(n=6)** as well as in the secondary heart field appending to the atrium of the heart tube (Supp. Figure 4). Notably, the tissue overlying the forming outflow tract, ventricle, and prospective atria is enriched in NO. (Fig 1I) NO signaling is also observed broadly across the forming vascular plexus and angiogenic tissues overlying the primitive heart tube (PHT) coincident with primitive angiogenic tissue. NO signaling here corresponds with its role in development of the nascent embryo vasculature (Huang *et al*., 2010). Furthermore, in later stages of development vascularizing tissue overlying the forming heart tube could be seen with high NO levels. (Sup. 6B). Interestingly, NO elevations are also noted in the intervening lateral plate mesoderm in between nascent somites (Figure 2 I-K). Our data demonstrate broad low levels of elevated NO and focal hot spots of high NO signals encompassing the ongoing anterior heart morphogenesis and foregut formation progression, suggesting indicating many functions of NO signaling in the early embryo development. Biological replicates of age matched embryos are depicted in Supplemental Figures 1–4.

**Figure 2.**
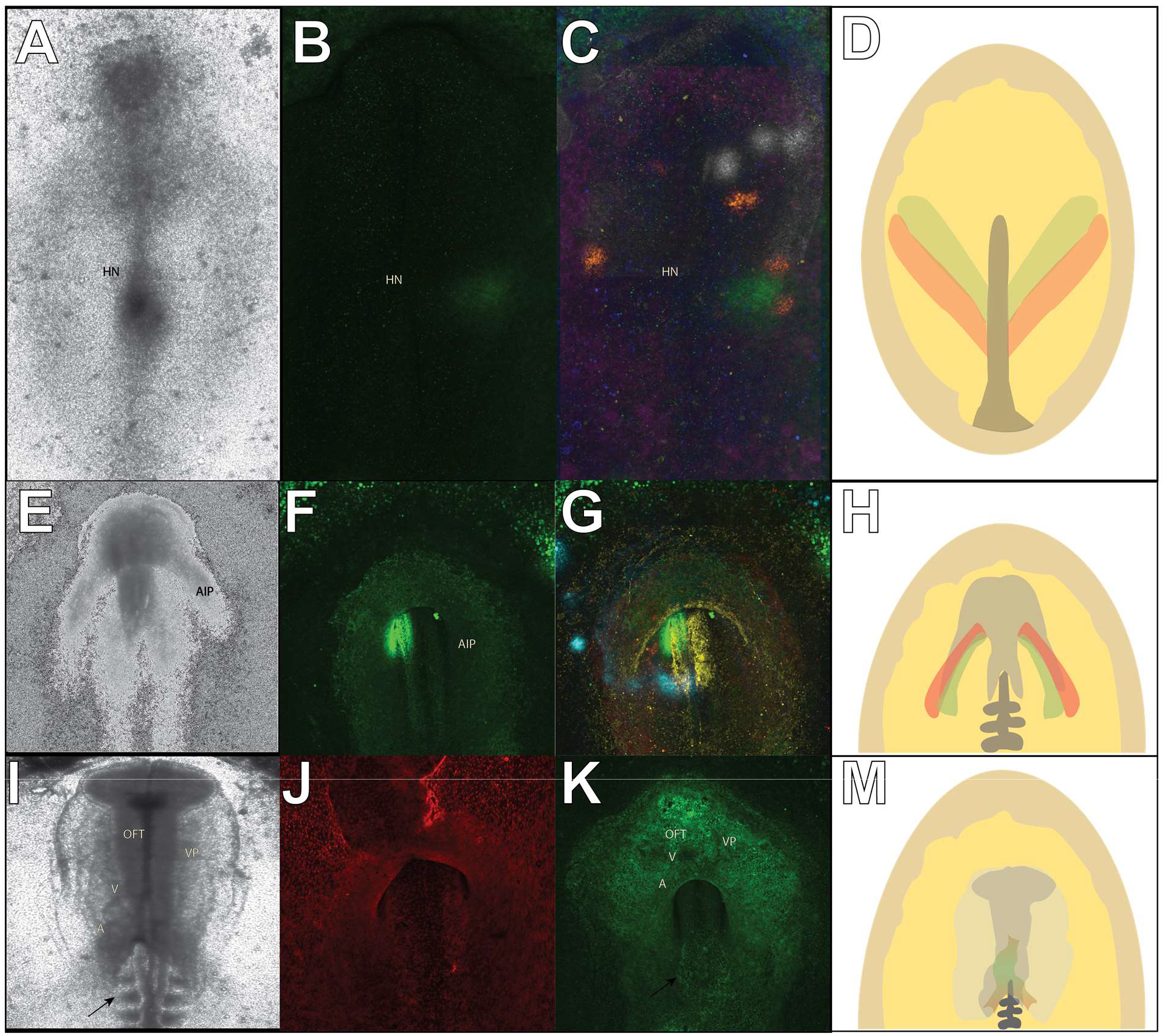
NO formation dynamically forms “hotspots” in correlation with the heart forming regions. Ventral view of transmitted light images of embryos at HH stages (A) 4-4+, (E) 6-7, and (I) 9-10 respectively. (D,H,M) Age matched cartoons with first heart field progenitors in red, and second heart field progenitors in green (B) HH4-HH5 staged embryos exhibit elevations of NO in the bilateral HFRs), as well as focal “hotspots” lateral to the extending Hensen’s Node (HN). However, a variety of patterns of NO formation were observed. (C) A composite image from 6 separate embryos demonstrate both diffuse elevations (blue, magenta, yellow) across the lateral endoderm (n=3), as well as focal elevations (green, orange, white) in both the left and right lateral endoderm (n=3). (F,G) HH6-7 embryos exhibited both broad, and NO signaling patterns in the head fold, cardiogenic splanchnic mesoderm, and the folding bilateral HFRs(yellow, red, green, & cyan). (J,K) NO signaling in HH9-10 embryos was diffuse across the ectoderm, but visibly enhanced in tissues overlying the fusing endocardial tubes and forming heart tube (J,K). Black arrow denotes the forming somites. (A-Atria, AIP-Anterior Intestinal Portal, OFT-Outflow Tract, V-Ventricle, VP-Vascular Plexus).

Our data demonstrates that at early stages of heart tube development, the endoderm produces NO to signal the cardiac splanchnopleure, but later monitoring of NO could not be easily performed as morphogenesis causes the embryo to make deep folds of cellular layers in the embryo head and heart regions from embryo morphogenetic growth. However, the overall pattern of NO formation in the pre-heart tube embryo coincides with cardiogenic and signaling tissues likes the lateral endoderm, cardiogenic mesoderm, splanchnic mesoderm, and nascent epicardium. NO signaling is highly dynamic, lending to variability of observed signaling patterns across embryos at similar points. However, we demonstrate elevations of NO coincident with the migrating cardiac progenitors and HFRs as they undergo morphogenesis and are signaled for cardiogenesis. Although dynamic NO signaling regulates numerous cellular processes, these observations are suggestive of endoderm NO paracrine signaling in gastrulating embryos to nascent cardiac progenitors in development of the Primitive Heart Tube (PHT).

### Endoderm NO Signals Differentiation of Myocardial and Endocardial Cells of the Primitive Heart Tube

We consider the possibility that NO signals in the cardiac forming regions of the embryo to enhance cardiac progenitor cells’ proliferation and differentiation throughout primitive heart tube development. We show endoderm NO hot spots in dynamic patterns in the cardiogenic development fields to support NO signaling in normal embryo heart formation (Figure 1). To investigate aberrant endoderm NO signaling in nascent cardiac progenitor cell differentiation and primitive heat tube formation, the endoderm layer was targeted with reagents to regulate NO formation by L-NAME to inhibit cNOS activity, or by donating NO directly through DETA-NONOate. These reagents respectively could reduce NO formation or elevate NO levels in the endoderm layer overlying the cardiac mesoderm (Sup 5 A-C). The NO reagents were diluted in Tyrodes saline solution and injected into the sub-germinal cavities of HH4-5 staged embryos, subsequent embryo growth and development to heart tube formation was monitored and effects on NO formation assessed with DAF2-DA (HH11). Embryos treated *in ovo* with 5mM L-NAME at HH4 for 1 hour, exhibited significantly decreased NO formation in ventral endoderm compared to control treated embryos, verifying this method for *in ovo* targeting of endoderm NO formation. (−39.9± 0.08%, n=6,p=0.01)(Sup 5C). Embryos exposed to higher levels of cNOS inhibition, L-NAME (25 mM) or greater resulted in severe abnormalities in heart looping, as well as profound defects in cranial morphogenesis and overall embryo size reduction. (Sup 6.A-C) These results demonstrate that a reduced NO signaling in the endoderm alters normal heart tube formation and looping, and induces severe defect in cranial development affecting head turning, optic vessel formation, and an overall reduced embryonic development depicted by truncated embryo morphogenesis and size.

We next determined the effects of NO signaling in endoderm on heart development by cNOS or NO donor treatments given at gastrulation stage (HH4-5), and monitored for myosin heavy chain expression in HH11/12 embryos. Results demonstrate that NO donor, 2μM DETA-NONOate(n=6), produced increased levels of MHC expression compared to MHC levels in vehicle-treated control (n=5) age-matched embryos. Conversely, cNOS inhibitor, 5mM L-NAME(n=7), treatment resulted in a reduced MHC expression (Fig 3 A-C). These visual differences in MHC expression in heart tubes demonstrate that NO signals in the nascent differentiation of cardiomyocytes. However, this result does not provide insight to the effects of NO signaling on the differentiation of the two cell types of the heart tube, myocardial and endocardial. We then assessed effects of NO signaling on differentiation of cardiac progenitors towards the two tissue types that constitute the majority of cells in the heart tube. To do so, embryos were treated with NO regulator at HH stages 4-5 as previously described, the embryos were then isolated, homogenized, and the protein lysates probed for endocardial marker, VE-Cadherin(VE-CAD), and MHC, marker for myocardial cell(Fig. 3D). The inhibition of NOS via 5mM L-NAME resulted in heart tubes expressing significantly less MHC (−37.4% ± 0.07%, n=7, p<0.000) compared to heart tubes from age-matched control (Fig3E). Conversely, NO donor, via 2uM DETA-NONOate, resulted in heart tubes expressing a greater amount of MHC (+14.4%± 0.03%, n=7, p<0.000) compared to heart tubes from age-matched control and that was significant (Fig. 3E). Furthermore, 2uM DETA-NONOate enhanced expression of VE-CAD in heart tubes (+27.6%±0.01%, n=3,p=0.03) compared to controls, but NOS inhibition did not have a significant effect on VE-CAD expression (+10.4%± 24.5%,p=0.3) (Fig. 3F). These results suggest elevated NO signaling in both endocardial and myocardial differentiation. When comparing ratio of myocardial to endocardial markers in the heart tube by normalizing VE-CAD expression to MHC, both NO donor & cNOS inhibitor treatments resulted in enhanced ratios compared to that of control heart tubes, (+88.2%±14.4%, n=3,p=0.01) & (+11.3%±0.03%, n=3,p=0.00) respectively (Fig. 3G). We demonstrate elevated endoderm NO signaling enhances expression of both endocardial and myocardial markers in heart tubes compared to controls, while increasing the ratio of myocardial to endocardial markers. Conversely, inhibition of NOS and reduced NO decreased expression of myocardial markers in heart tubes and also resulted in an increase in the ratio of endocardial to myocardial cells. These results show clear NO signaling roles in the augmentation of proliferation and differentiation of cardiac progenitor cells in HFRs through effects on endocardial and myocardial development. Although the relationship between NO signaling and the ratio of endocardial and myocardial cells in the heart tube is unclear, future work will investigate NO signaling on endocardial and myocardial cell proliferation at this early time in heart tube development. Our findings here demonstrate chicken embryo cardiogenesis is under NO signaling regulation, and for the first time, we show consistent low level elevated NO and dynamic NO signaling hot spot in the endoderm layer in correlated, close association with cardiac mesoderm, to suggest active NO paracrine signaling in the developing heart tube.

**Figure 3:**
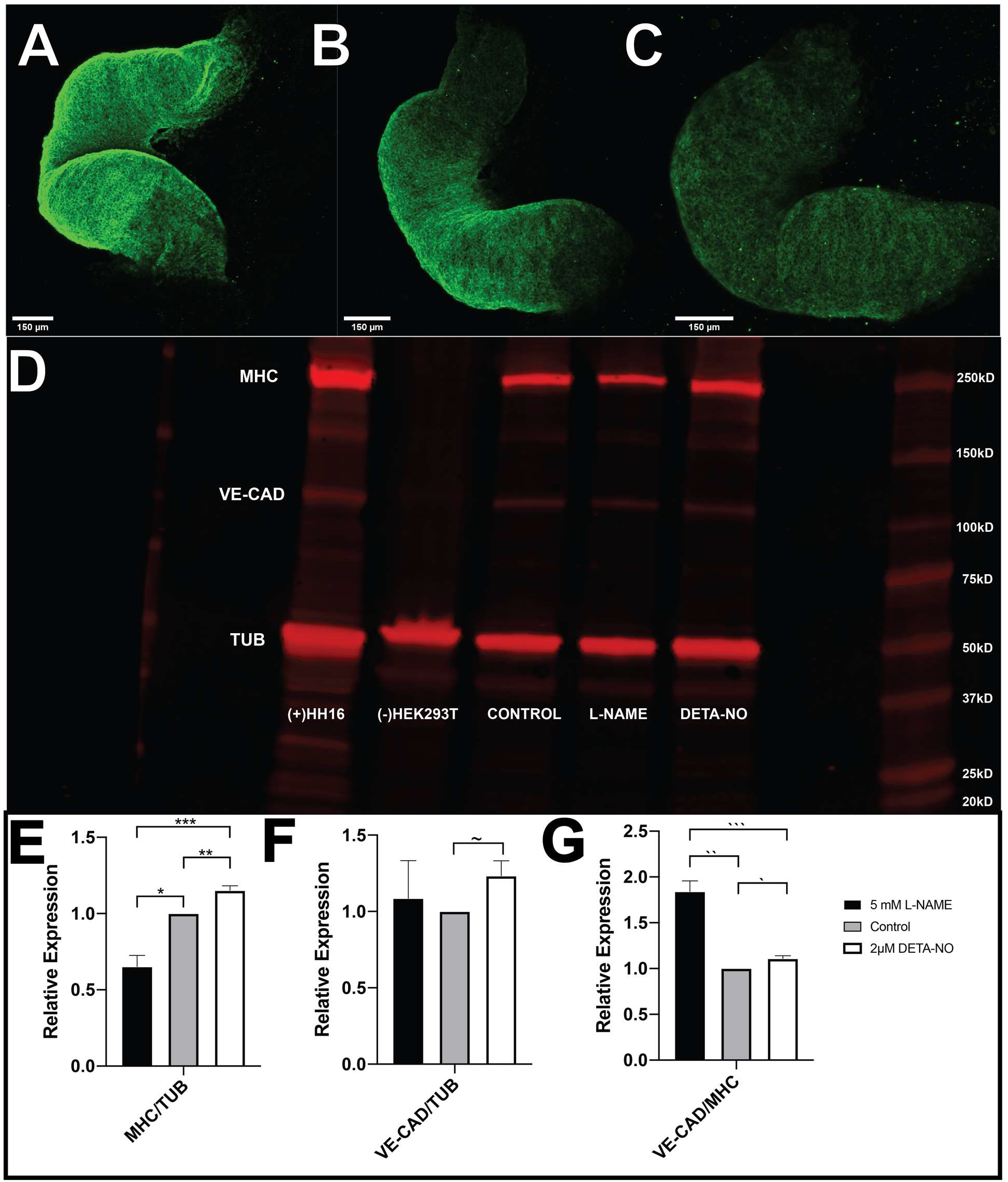
Endoderm NO enhances expression of endocardial and myocardial markers in the primitive heart tube. Embryos endoderm were treated with either (A) 5 mM LNAME, (B) vehicle Tyrode’s Saline, or (C) 2uM DETA-NONOate at HH5-6. Primitive heart tubes (PHTs) were stained for embryonic myosin heavy chain 1<u>°</u>, and Alexa 488 2<u>°</u>(maximum Z-projections of 30 stacks with 10 uM spacing). (D) Western blot of total protein from PHTs were probed for vascular-endothelial cadherin (VE-CAD) (115 kDa), myosin heavy chain (MHC) (220kDa), and tubulin (TUB) (55kDa) as control. (E-F) Quantitative analysis of MHC and VE-CAD expression was performed as normalized to TUB and (G) VE-CAD normalized to MHC as a comparison of endocardial to myocardial markers (n=3) in PHT total protein fractions. Exogenously donated NO resulted in a significant increase in MHC and VE-CAD expression compared to both control treated embryos (P<0.00,n=7). Conversely, inhibition of NO signaling significantly attenuated MHC expression compared to control treated embryos(P<0.01). one-tail Student’s t-test (***P<0.00002, **P<0.0001, *P<0.0003) (~P<0.04)(***P<0.002, **P<0.001, *P<0.01).

### NO regulates cardiomyocyte proliferation

NO is a potent regulator of proliferation, with high NO levels (uM) signaling for cell cycle arrest/senescence and low NO levels (pM-nM) for proliferation. (Thomas *et. al.* 2009). We hypothesize that low level NO signaling in the endoderm may signal to regulate proliferation of cardiac progenitors in the myocardium of the heart tube. In numerous cell culture systems, NO signaling has been implicated in regulating proliferation and differentiation activities with low NO levels signaling in proliferation, and high NO levels signaling in cell cycle arrest, cellular differentiation or senescence (Gooch *et. al.* 1997, Kanno *et. al.* 1999, Kosonen *et. al.* 1997). As the previous experiments demonstrated an increase in myocardial markers in response to NO, we hypothesized the paracrine produced endoderm NO might signal to maintain or augment proliferation of myocardial cells in the hear tube. To investigate NO in the proliferation of myocardial cells in the heart tube embryos were subjected to overnight NO reagent treatment at HH 4-5 as previously described, and subsequently ventrally and dorsally pulsed with 100 ul of 500 uM Edu diluted in either 5 mM L-NAME, 2 uM DETA-NO, or control, Tyrode’s Saline solution, at heart tube formation (HH10), and then incubated for 4 hours to allow for incorporation of the Edu thymidine analog. (Warren, Puskarczyk and Chapman, 2009) Embryos were then imaged, and the proliferative, Edu+, and total nuclei, Hoescht+, cells were counted. For this task, we have developed an automated method for quantifying the number of nuclei, as a manual counting of nuclei in concordance with volume of images was infeasible. Images were stitched and analyzed using Fiji (Preibisch, Saalfeld, and Tomancak, 2009). The cell counting is automated by training a U-net style autoencoder that is trained to predict high scores near the centroid of each chicken nucleus in a 2D image. We have observed that the max-projection of nuclei does not result into serious overlap and occlusion because of their typical distribution in 3D space (Figure 4). Thus, we have done all our computation in the max-projected 2D space instead of the 3D. In fact, to keep the dimensionality of our data small we have further split the 512 × 512 max-projection into four equal image tiles for the autoencoder to process. During inference, each test image is split into four equal sized image tiles as shown in Figure 3 in the supplement. The output scoremaps of the four tiles are collected and placed in the appropriate grid position to build the prediction scoremap for the full image. The predicted scoremaps and the subsequent peak detection deliver the final coordinates of the nuclei centers. The estimated nuclei centers are compared with the ground truth to ascertain the effectiveness of the methodology we have followed in this work. With this methodology we have been able to achieve 71% precision and 69% recall with an F-Score of 70%. Of course, the neural net makes errors especially in densely populated areas, but even then, our biologist has to work reasonably less in correcting its prediction mistakes as compared to the situation when he/she would have to annotate from scratch. Once evaluation is done and we have fixed the choice of our parameters, we train a final neural net using the entire training set, in order to use this trained model for predicting the nuclei positions in all of the unseen image sets. This step automates the long and slow process of manually identifying each and every nuclei in the large image dataset from scratch. We then counted the nuclei residing in the ventricular myocardium, as this tissue is the first to develop terminally differentiated cardiomyocytes in the primitive heart tube (Fig.5 A,E,I). As individual nuclei did not dual stain for Hoescht and Edu, proliferation was calculated by taking Edu+ as a percentage of the sum of Edu+ and Hoescht+ cells. NOS inhibition (Fig.6 A-D) significantly attenuated proliferation of ventricular myocardial cells by 7.8% ± 3.3%,(p=0.04,n=8), in comparison to vehicle treated embryos(Fig 5 E-H). Conversely, donation of endodermal NO (Fig. 5 I-L) resulted in a 7.6% ± 5.6%,(p=0.09,n=5) increase in proliferation in comparison to control-treated embryos. (Fig. 5M) Overall, we observed that NO signaling demonstrates regulatory effects on ventricular cardiomyocyte proliferation by sustaining normal proliferation. However, these effects on proliferation are not acute, suggesting that endodermal NO potently regulates terminal differentiation of cardiac progenitors in formation of the heart tube while marginally regulating proliferation of myocardial cells.

**Figure 4.**
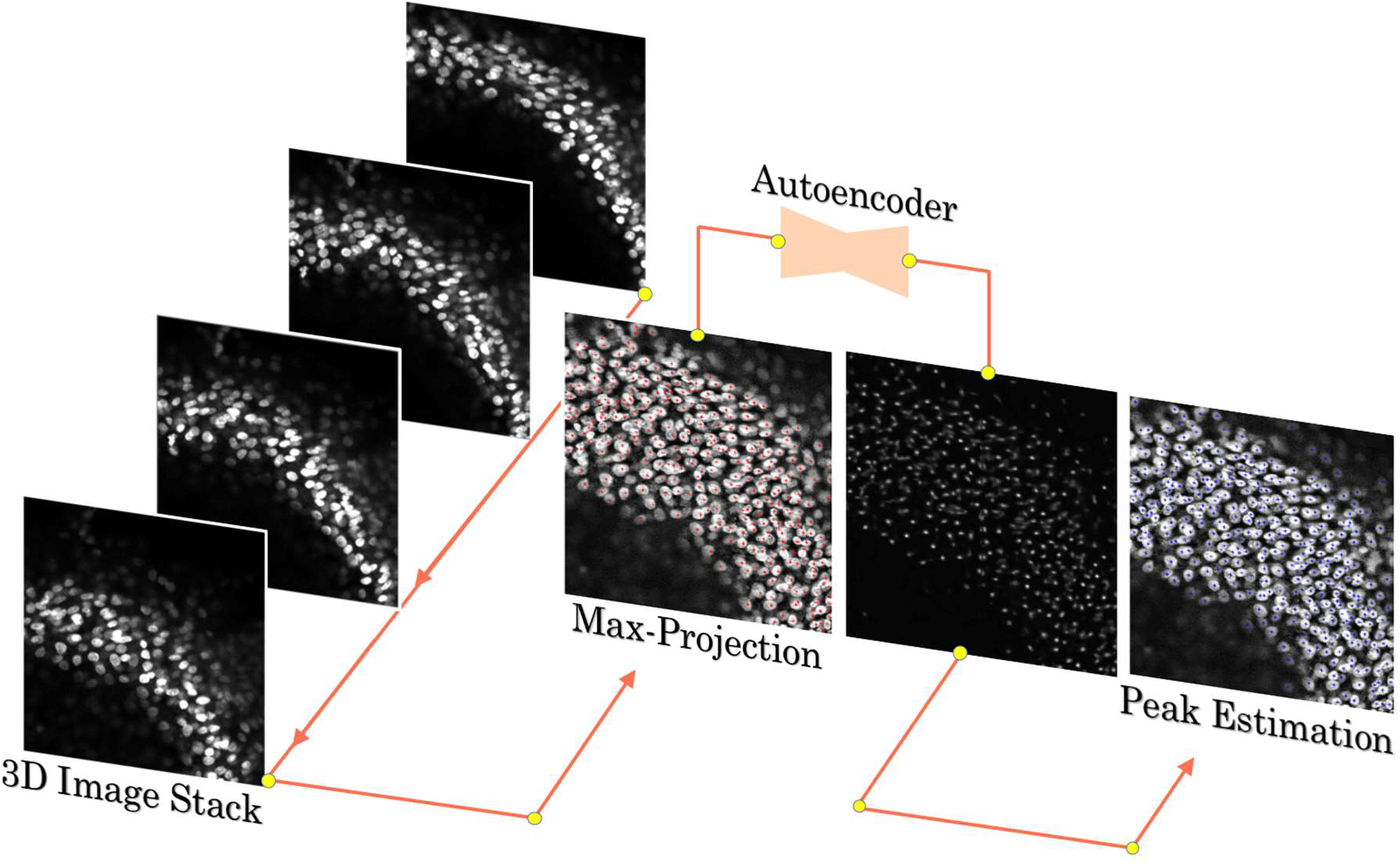
Methodology for high dimensional matrix regression task. We derive a 2D max-projected image from the 3D stack of confocal slices. Note the shifting pattern of the nuclei in the z-slices avoid any major occlusion/overlap in the max-projected image. We train our deep neural autoencoder with the max-projected images to identify the positions of the cell nuclei. In response to a typical 2D max-projected image as shown in the figure, the deep autoencoder predicts gaussian peaks as a result of high dimensional matrix regression. A 2D peak finding algorithm locates the gaussian peaks in order to predict the center locations of the nuclei. Please see details in the supplementary text.

**Figure 5.**
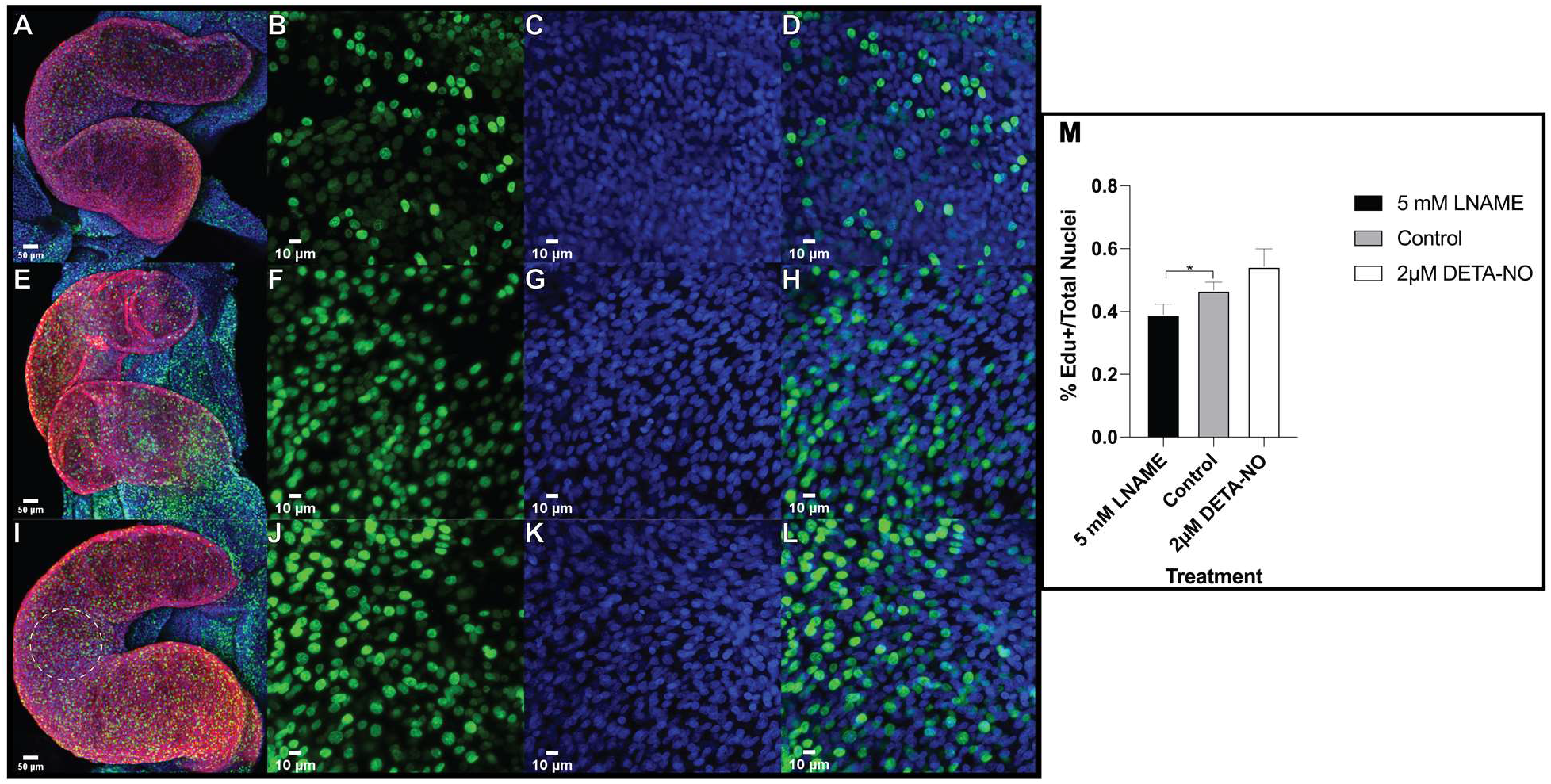
NO signals to maintain proliferation in the myocardium of PHT. (A, E, I) PHT of embryos aged HH12 (n=3) triple stained for MHC(red), proliferative Edu+ cells (green), and total nuclei Hoescht+ (blue) (maximum Z-projections of 30 stacks with 1 uM spacing) High magnification (40X) images of the (B, F, J) Edu+, (C, G, H) Hoescht + and (D, H, L) merged channels of the ventricle myocardium. Dashed white circle indicates the region from which high magnification images of the ventricle were taken and used for quantification of proliferative and total nuclei. Nuclei were counted as described in Figure 4. And supplementary text. (M) Total (Hoescht+) and actively proliferation (Edu+) cells were quantified in the left ventricles of PHTs treated with 2 uM DETANONOate, 5 mM-LNAME, and vehicle and compared. NOS inhibition significantly impaired proliferation of cells in the ventricular myocardium (P<0.04). 1 tailed student’s T-test. (*P<0.04)

**Figure 6.**
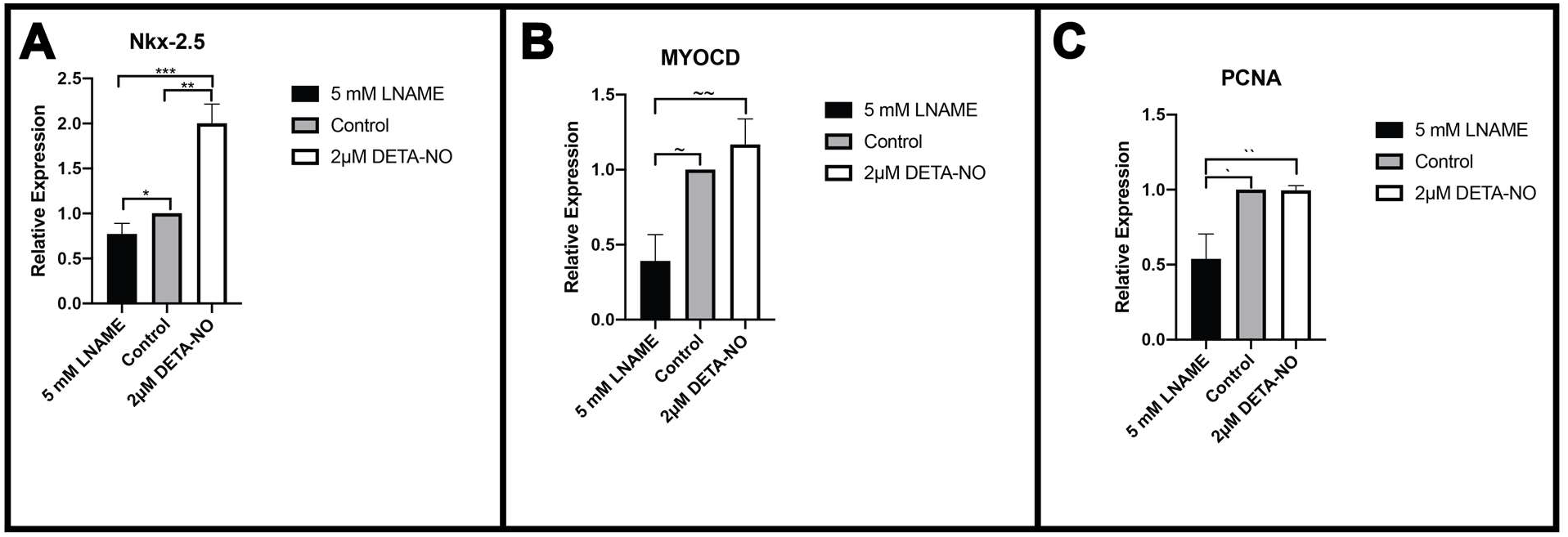
Endodermal NO regulates expression of key cardiogenic transcription factors. qRT-PCR analysis of key cardiogenic transcription factors, (A) *Nkx-2.5* and (B) myocardin and (C) PCNA normalized to housekeeping gene GAPDH. Exogenous NO significantly enhanced expression of *Nkx-2.5* in comparison to control (P<0.001), but did not have significant effects on *MYOCD* or PCNA. Conversely treatment with LNAME significantly reduced expression of Nkx-2.5, *MYOCD*, and PCNA in comparison to control and DETA-NONOate. One tailed student’s T-test. (A) (***P<0.0005, **P<0.0008,*P<0.04), (B) (~~P<0.01, ~P<0.004), (C) (**P<0.01, *P<0.02)

### Cardiogenic genes regulated by NO formation

Our previous experiments demonstrate regulation of myocardial differentiation and proliferation by endodermal NO, but effects of NO signaling on expression of key cardiogenic transcription factors throughout gastrulation is still unknown. *In vitro* experiments have demonstrated NO signaling to regulate expression of nascent cardiac transcription factors for developing cells in the cardiac muscle cell lineage (Mujoo *et al*. 2006 & 2008, Tejedo *et al*., 2010). To assess endoderm NO signaling effects on nascent cardiac progenitor gene expression, we treated HH5 staged embryos with NO reagents for 4-6 hr *in ovo* culture, explanted endocardial tubes and surrounding tissue(HH8), and quantified levels of early cardiogenic transcription factors *Nkx-2.5* and *myocardin(MYOCD)*, as well as proliferating cell nuclear antigen*(PCNA)*. In response to NOS inhibition, expression of both cardiac transcription factors were significantly attenuated, with *Nkx-2.5* and *MYOCD* reduced −22.5% ± 11.5%,(p=0.04,n=5) and −60.8 ± 17.5%, (n=4, p.004) respectively, compared to control tissues. (Figure 6. A-B) Surprisingly, inhibited NO signaling also significantly reduced proliferation, −46.2% ± 16.6%,(n=4, p=.01). (Figure 6. C) Conversely, elevated NO signaling resulted in increased expression of both *MYOCD* and *Nkx-2.5*, +16.8% ± 17.0% (n=4,p=0.18)& 100.2% ± 21.3%(n=5,p=0.00) respectively, but only *Nkx-2.5* expression significantly impacted by increased NO signaling. Oddly, enhanced NO formation did not significantly impact proliferation, resulting in a −0.004% ±0.031%(n=5,p=0.4) reduction in PCNA expression. Overall, these experiments demonstrate a pivotal role for NO signaling from endoderm in development of nascent cardiomyocytes through regulation of key cardiac transcription factors, *Nkx-2.5* & *MYOCD*.

## DISCUSSION

At the earliest development stages, mouse embryos produced NO under estrogen regulation to facilitate uterus implantation and normal development of the embryo (Gouge *et al*., 1998) thereby, showing the importance of the NO messenger molecule to signal in nascent embryo development. *In vitro* cultures of human and mouse ESC also demonstrated NO signaling ability to promote cardiogenesis, as shown by changes in ESCs in cardiac gene expression and cardiomyocyte proteins to support differentiation into cells in the cardiac lineage. (Kim *et al*., 2004; Mujoo, Krumenacker and Wada, 2006; Mujoo *et al*., 2008; Pelaez *et al*., 2017) The heart is the first organ formed in the embryo and we sought to determine if NO signals in early embryo heart formation and to asses NO regulation of bilateral HF cardiogenesis. Here, we report for the first time, that the during gastrulation the endoderm layer dynamically forms the NO messenger to autocrine and paracrine signal in spatiotemporal patterns that are coincident with nascent cardiogenesis in the bilateral heart fields (HFs) through heart tube formation. Through *in vivo* manipulation of NO production in the endoderm layer adjacent to the bilateral HFs, we demonstrate NO signaling enhances differentiation of cardiac progenitor cells into cardiomyocytes through an effect on cardiogenic transcription factors *Nkx-2.5* & *myocardin*. We also demonstrate NO signaling regulation over heart cell lineage development by immunolabeling of the cardiomyocyte marker, myosin heavy chain(MHC), and the endocardial cell marker, vascular endothelial cadherin *(VE-CAD)*. Furthermore, we demonstrate that endoderm NO maintains proliferation of cells by quantification of actively proliferating cardiomyocyte nuclei labeled with thymidine analog, EdU (5-ethynyl-2’-deoxyuridine), as well as transcription of proliferative marker, PCNA. Finally, acute alterations in endoderm NO signaling levels result in heart tube formation defects as well as abnormalities in cranial structures, suggesting a multifaceted role for NO signaling in nascent embryonic development. These findings suggest a model of endoderm NOS mediated NO formation and autocrine and paracrine signaling to support cardiogenic development in bilateral HFs and in heart tube formation (Fig 7). We report a novel finding in embryo cardiogenesis that the endoderm layer produces significant amounts of the NO messenger molecule to paracrine signal into cardiac mesoderm to enhance differentiation and proliferation of cardiac progenitors in the developing HFs.

**Figure 7:**
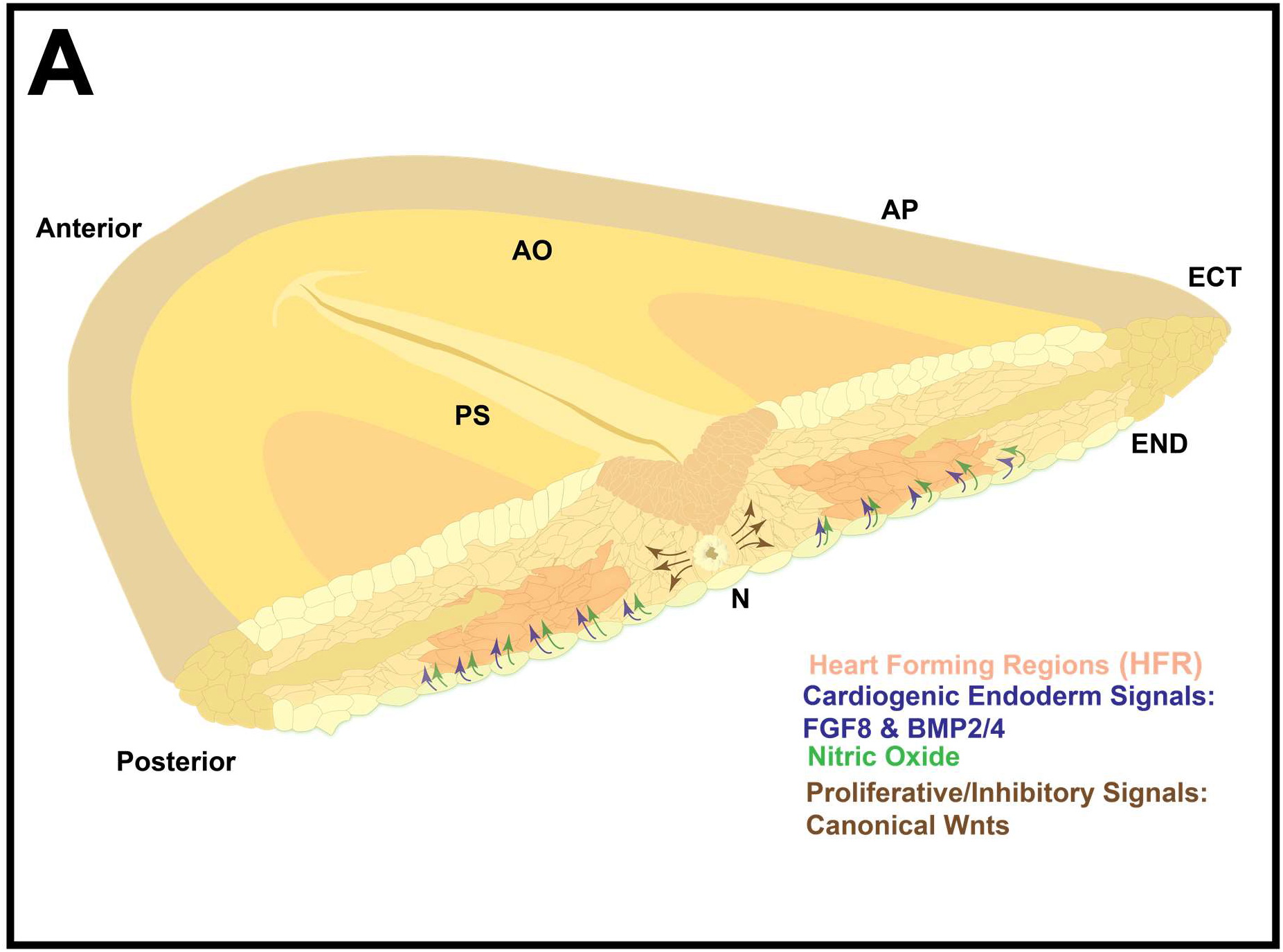
Endoderm NO signals in concert with morphogens to regulate cardiogenesis. (A) Representative pictograph of NO signaling and related morphogen signaling in the developing heart fieldto suggest NO generated in the endoderm (green) may support cardiogenic development similar to other cardiogenic endoderm signals such as FGF and BMP2/4 (blue) together with the signaling activities of the surrounding tissues like the notochord (N). (AO - Area Opaqua, AP-Area Pellucida, END-Endoderm, ECT-Ectoderm, PS-Primitive Streak)

Our data shows that endodermal NO signaling enhances the expression of cardiogenic transcription factors *Nkx-2.5* and *myocardin* in the forming endocardial tubes (Fig. 6). This result is consistent with *in vitro* studies which demonstrate an excess of NO induces expression of cardiogenic transcription factors including *Nkx-2.5* and differentiates cardiac progenitors to cardiomyocytes. (Kim *et al*., 2004; Nikonoff *et al*., 2008) Furthermore, inhibition of NO signaling in cell cultures of human and mouse ESCs results in a reduced expression of key cardiac transcription factors, *Nkx-2.5* and *Gata4*, as well as a decrease in cardiomyocyte proliferation (Mujoo, Krumenacker and Wada, 2006; Mujoo *et al*., 2008; Lopez-Sanchez *et al*., 2009; Tejedo *et al*., 2010; López-Sánchez and García-Martínez, 2011; Jankowski *et al*., 2012). *Myocardin* is one of the earliest cardiac transcription factors expressed in the HF (Warkman *et al*., 2008), with our data we show it is also regulated by NO. *Myocardin* also acts as a transcriptional co-activator of the cardiac transcriptional program by enhancing expression of *Nkx-2.5*, suggesting that NO by activation of its canonical sGC/cGMP pathway may play a central role in supporting the maintenance and differentiation of cardiac progenitor cells for later heart tube development (Wang *et al*., 2001). NO canonical signaling, through cGMP production and activation of different protein effectors including cGMP-dependent PKG1, has been shown to activate BMP4, (Traister *et al*., 2004; Schwappacher *et al*., 2009) itself capable of inducing Myocardin and *Nkx-2.5* transcriptional activation via Smad1/5/8 signaling(Wang *et al*., 2012). Our work in the chicken embryo which shows an ability of NO signaling to upregulate early cardiac transcription factors, myocardin and *Nkx-2.5*, in the developing HFs, is consistent with previous findings. Furthermore, previous work has demonstrated that transient NO production enhances differentiation of mouse embryonic stem cells *in vitro* by inducing nuclear translocation of the key cardiogenic transcription factor, cardiotropin-1, to induce expression of numerous cardiac transcription factors and cardiomyocyte protein α-actinin, necessary for induction of cardiomyogenesis in embryonic stem cells.(Mascheck *et al*., 2015) Though our work did not investigate effects of NO on *cardiotropin*-1, *GATA4*, or other early cardiac transcription factors, endoderm NO signaling may also regulate activity of these early cardiac transcription factors for *in ovo* HF development, suggested by results from *in vitro* studies (Friart *et al*., 2016). In terms of differentiation and cardiomyocyte maturation, NO regulation over expression of cardiomyocyte contractile proteins has been demonstrated in mouse embryonic stem cells cultured with NO donor compounds, like DETA-NONOate, to produce/enhance functional cardiomyocyte differentiation shown through the expression of cardiac muscle proteins, like cardiac troponin and myosin heavy chain (Kim *et al*., 2004; Friart *et al*., 2016). Our data for *in ovo* cardiomyocyte development is consistent with these *in vitro results*, demonstrating that NO signaling reductions by NOS inhibition cause a marked reduction in cardiomyocyte expression of *MHC*, and endocardial cell marker, *VE-CAD*, while elevated NO signaling enhanced expression of early cardiac transcription factors as well as cardiomyocyte MHC protein. We also assessed if this effect on enhanced cardiac transcription factor and cardiomyocyte protein expression was caused by NO signaling ability to enhance cellular proliferation, as NO regulatory activity in proliferation has been shown in other systems.

NO signals to regulate proliferation in various tissues (Kosonen *et al*., 1997; Traister *et al*., 2004), and in our experiments NO signaling maintains proliferation of the cardiac progenitor cell pool in the bilateral heart fields to produce the heart tube left ventricle that derives from first HF cardiac progenitors and cardiomyocytes. This is consistent with a study that demonstrated that through eNOS activation, NO canonical signaling is necessary for proliferation and differentiation of neonatal cardiomyocytes *in vitro* embryoid culture, and *in vivo* postnatal heart development.(Lepic *et al*., 2006) However, NO signaling has differential effects on proliferation in various cell types indicative of complex regulatory actions of canonical NO signaling that is not well understood. For example in the developing neural tube, NOS inhibition resulted in greater proliferation while excessive NO levels via NO donors, reduced cell proliferation (Traister *et al*., 2004). Additionally, *in vitro* culture of adult rat aortic smooth muscle cells NO canonical signaling signal, via raising intracellular levels of cGMP, has been demonstrated to inhibit proliferation by inhibiting thymidine incorporation during DNA synthesis. (Garg and Hassid, 1989) Though our data and previous studies demonstrate a causal relationship between NO levels in signaling for proliferation, the mechanisms by which NO signaling elicits proliferative responses in some cell types like cardiac progenitors, but inhibits proliferation in other cell types, like vascular smooth muscle cells, are not well understood but likely influenced by interactions with regulatory morphogens associated with cell proliferation.

Cardiogenic inhibitory morphogen signaling like the non-canonical *Wnts* 3c and *Wnt* 11, and *Sonic Hedgehog (SHh)* are produced along the embryo midline in neural tube, notochord and in the adjacent splanchnic mesoderm, maintaining cardiac progenitors in these mesodermal regions in proliferative, pre-cardiogenic states (Marvin *et al*., 2001b; Foley and Mercola, 2005; Tzahor, 2007; Kelly, Buckingham and Moorman, 2014). We show that NO formation was diminished along the embryonic midline where inhibitory signals to delay differentiation and cardiogenesis is repressed (Marvin *et al*., 2001b; Cambier *et al*., 2014). This observation may be a result of bidirectional cross-regulation of NO signaling with canonical Wnts, where β-catenin associated eNOS inhibits formation of active, phosphorylated eNOS, to mediate low NO production. (Passacquale *et al*., 2014). However, canonical NO signaling may also accentuate downstream Wnt signaling via inhibition of *GSK3β* to allow for nuclear translocation of β-catenin and transcription of its downstream targets. This relationship may contribute to the pattern of observed NO signaling where NO formation is enhanced in the pro-cardiogenic signaling tissue but diminished in cardiogenic inhibitory tissues like the embryo midline, as it is known that NO and canonical Wnts demonstrate cross-regulatory action in initiation of skeletal muscle regeneration (Kim *et al*., 2004; Drenning *et al*., 2008) The NO messenger molecule pleiotropically signals in the body in every organ system in normal physiology and pathophysiology conditions. (Moncada, Palmer and Higgs, 1991; Marshburn *et al*., 2005; Martínez-Ruiz, Cadenas and Lamas, 2011) However, investigation into NO signaling in embryogenesis are relatively new. We show here for the first time, chicken embryo endoderm NO paracrine signaling in dynamic patterns to correlate with pre-cardiac mesoderm throughout early gastrulation in the cardiogenic regions of the chicken embryo, as well as effects of this signaling activity in regulating the differentiation and proliferation of nascent cardiomyocytes in the cardiac mesoderm of the first and secondary heart field during gastrulation through heart tube formation. Our observed effects of manipulation of endoderm NO signaling on cardiac progenitor development is consistent with NO signaling effects in previous studies which found NO signaling to regulate differentiation and proliferation of cardiac progenitor cells *in vitro* (Kim *et al*., 2004; Mujoo, Krumenacker and Wada, 2006; Tejedo *et al*., 2010; Friart *et al*., 2016), as well as *in vivo* later embryonic development of the multichambered heart. (Bloch *et al*., 1999; Lepic *et al*., 2006; Liu and Feng, 2012). Collectively these experiments demonstrate that endoderm NO signals to regulate cardiogenesis *in ovo* at the earliest timepoints of embryonic heart development during the specification of cardiac progenitor cells for heart formation.

## Acknowledgements

We thank Lisa Galli for providing pivotal guidance in experimental procedures, as well as generously providing antibodies and laboratory tools used in this study. We also thank Dr. Takashi Mikawa for constructive discussions and his expertise in the field of embryonic cardiology. We thank Annette Chan and the Cell Molecular Imaging Center at SFSU for technical support with microscopy. Y.-H. M. C. and W. L. C. acknowledge funding from NIH award 1SC2GM118267 and the NSF STC Center for Cellular Construction award DBI-1548297. S.B. and S. K. B also thankfully acknowledge the NSF STC Center for Cellular Construction award DBI-1548297.

## Methods

### 1.1 Animal Procedures

Fertile White leghorn chicken eggs from Skippy’s Eggs Store, Petaluma, CA, were stored at 14°C no longer than two weeks, before being incubated at 39 °C at ~90% humidity. All tools used for embryo dissection and manipulation were sonicated in DI water and glass bead sterilized before use.

### 1.2 Short-Term Culture for Live Imaging

Embryos were isolated and cultured using modifications to New’s whole embryo culture technique <u>(New 1955; Chapman *et al*. 2001)</u>. Cultures were labeled with 100 uL of 5uM DAF-2DA in a humidified chamber for 1 hour before embedded in 2% low melting agarose for fluorescence confocal microscopy.

### 2.1 Pharmacological Endoderm treatment & Embryo Isolation

Eggs were incubated 20-22 hours for HH5-6 embryos. 100uL of DETA-NONOate **(Cayman Chemical, Item No. 82120)**, L-NAME**(Cayman Chemical, Item No. 80210)**, and or sterile Tyrode’s **(Sigma-Aldrich, T2145)** were injected *In Ovo* into the subgerminal cavity. Eggs were resealed with parafilm and placed back into the incubator until the desired stages were reached. Embryos were subsequently extracted, washed, and preserved in PBS on ice for following downstream applications.

### 2.2 Immunohistochemistry

Embryos were fixed for 1 hour at room temperature with 4% paraformaldehyde Embryos were blocked and permeabilized overnight with 4 % goat serum and 0.3% Triton-X**(Sigma, 9002-93-1)** in PBS. Embryos were labeled with 5% MF20 **(DSHB, frozen cells catalog#MF 20)** primary antibody in **diluted blocking solution** (2% goat serum and 0.15% Triton-X). Embryos were then treated with secondary antibody 0.25% Alexa Fluor 488, or 687, goat anti-mouse in **diluted blocking solution** overnight at 4 °C. Embryos were then placed in PBS at 4 °C for whole-mount imaging.

### 2.3 PHT extraction and Protein Isolation

HH11-12 heart tubes were cut with Vannes scissors at the omphalomesenteric vein*(OMV)* and the anterior outflow tract *(OF)*. Isolated heart tubes were then collected into a microcentrifuge tube with PBS on ice. Three hearts per condition were transferred into a lysis matrix tube (Bio 101, Inc. Lysing Matrix Cat. 6901 017) for centrifugation at 12,000 RPM in 4 °C for 3 minutes. Pellet was resuspended in 160ul of cold TENT lysis buffer (20mM Tris, 2mM EDTA, 150mM NaCl, and 0.25% Triton-X), a matrix lysing bead (Fisher, #MP116923050) was added to the tube and briefly vortexed. The lysate was transferred into a fresh tube containing 40 ul of 5x Laemmli Buffer. The sample was then fastened into a **Beaker Buddy Boiling Rack**, and placed into a 500 ml beaker containing boiling water over a Bunsen Burner, to boil for 5 min. The sample was allowed to cool then stored at −20°C for downstream gel electrophoresis and western blot. Negative Control, Hek293T cell lysates **(ATCC, CRL-11268)** were a gift from the Burrus Lab **(Lisa Galli)**.

### 2.4 Gel Electrophoresis and Western Blot

10 ul of protein standard (Odyssey® One-Color Protein Molecular Weight Marker) and 15 ul of each sample was loaded into a 4-20% gradient polyacrylamide gel (Mini-PROTEAN® TGX ™ Precast Gels). Sample lanes were flanked by lanes loaded with 15 ul of 1X Laemmli Buffer in TENT lysis buffer. Protein separation was performed in a biorad electrophoresis tank (Model 422 Electro-Eluter) filled with a running buffer containing 14.4 g Glycine, 3 g TRIS, and 0.1% SDS in 1L ddH2O. Proteins were then wet-transferred to a PVDF membrane with a running buffer containing **200 mL Methanol, 1.53 ml Ethanolamine, and 0.1 % SDS in 1L ddH2O.** Membranes were blocked with 10 mls of Odyssey Blocking Buffer **(Licor, 927-40000)** for 1 hour at room temperature before incubation with 3 different primary antibodies: 1:10 MF20, 1:4000 tubulin **(Santa Cruz Biotech, SC5274)**, and 1:3000 VE-CAD **(ThermoFisher Scientific, SC5274)** overnight at 4C. Blots were then washed 4 times with wash buffer 0.1% Tween20 (Fisher, BP337-100) in PBS and subsequently incubated in secondary antibody **(Fisher, A-21057)** for 2 hours at room temperature. Membranes were then washed 4 times with PBS+0.1% Tween20 and dried overnight. Secondary antibody fluorescence was detected using the Li-COR Odyssey Clx Imaging System. Fluorescence-based quantification of relative protein concentrations was then assessed using Image Studio - Odyssey Analysis Software and normalized to levels of tubulin, or MHC for VE-CAD/MHC comparison.

### 3.1 Edu Treatment and Immunostaining

Edu was diluted in 5 mM L-NAME, 2uM DETA-NONOate, and or sterile Tyrode’s solution (Warren *et al*. 2009). 100ul of 500uM Edu+NO reagent solution was injected into the subgerminal cavity and gently pipetted over the embryo’s dorsal side. Eggs were resealed and returned to the incubator for 4 hours. Embryos were subsequently harvested, fixed, blocked, and permeabilized as previously described. Embryos were then incubated in Click-iT™ Reaction cocktail (Invitrogen™ Molecular Probes™ Click-iT™ EdU Alexa Fluor™ 488 Imaging Kit) at room temperature for 1 hour. Embryos were rinsed three times before treated with 1:1000 diluted Hoescht 3342 overnight at 4C.

### 4.1 Cardiac Tissue isolation for RNA Isolation

The endocardial tubes and surrounding tissue was trimmed from HH8 embryos using Vannes scissors and fine tweezers. The tissue was then pipette transferred to a 1.5 ml snap-cap RNAse free microcentrifuge test tube on ice. The samples sat on ice for no more than 40 minutes before downstream RNA isolation.

### 4.2 RNA Isolation

Each sample prep for RNA isolation contained tissue from 3 embryos and was isolated using the Aurum™ Total RNA Fatty and Fibrous Tissue Kit. Quality of the eluted RNA was then assessed using the Nanodrop (product #). Only RNA samples with at least 100 ng/uL of RNA and a 260/280 ratio greater than 2.0 were used for cDNA synthesis and qRT-PCR

### 4.3 cDNA Synthesis

The Bio-Rad iScript™ cDNA synthesis kit was used for two-step reverse transcription quantitative PCR(RT-qPCR). The cDNA reaction was prepared with qPCR grade H2O with the total RNA sample at 25 ng/ul. The cDNA reactions were prepared in 200 ul RNase and DNase free PCR tubes. The tubes were then placed into a Bio-rad S1000 thermal cycler to be primed for 5 min at 25 °C following a reverse transcription step for 20 min at 46 °C, RT inactivation for 1 min at 95 °C, and left at 4 °C. The PCR tubes were then removed from the thermocycler, and 25ng/ul of cDNA were frozen for RT-qPCR. All experiments were conducted with -Reverse Transcription(RT) and -Template(Temp) controls, along with sample reactions, control(Tyrode’s), 5mM LNAME, and 2uM DETA-NO.

### 4.4 RT-qPCR assay

All qPCR reactions were prepared with SsoAdvanced Universal SYBR Green® Supermix, PrimePCR™ SYBR® Green Assay primers, and qPCR H2O. I All primers were analyzed using a CFX96 Touch™. qPCR was carried out for denaturation at 95 °C for 30 s following activation at 95 °C for 10 s, and annealing at 60 °C for 30 s for 41 cycles. Gene expression analysis was conducted using the Bio-Rad CFX Maestro™ Software for CFX Real-Time PCR Instruments software. The specificity of amplified transcript and absence of primer-dimers was confirmed by a melting curve analysis. Gene expression was normalized to GAPDH as an internal control. Relative mRNA expression was calculated using the comparative cycle threshold method (2-ΔCt). Information for the mixed forward and reverse Bio-Rad primers, specific for *Gallus Gallus*, used for the assay are described in the following table.

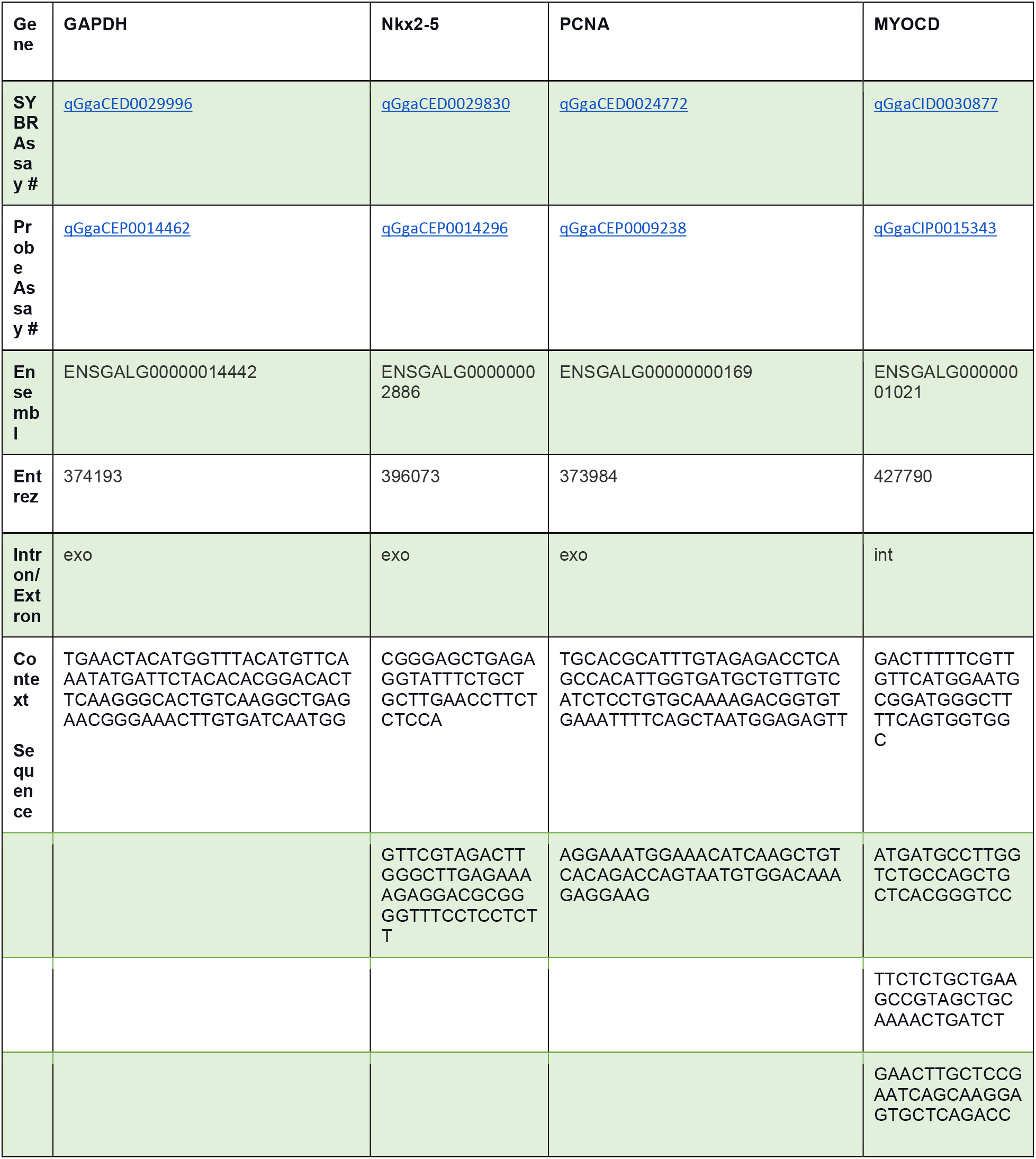

### 5.1 Fluorescence Confocal Microscopy

Confocal imaging of agarose embedded and whole-mount embryo specimens were performed using the Zeiss LSM 710 microscope and Nikon Plan Apochromat 5×, 10×, 20×, and 40xW objectives. Embryos labeled with DAF-2DA were imaged with 5× and 10× objective and with at least 25 slices at 10 uM spacing. Images of the whole PHT stained for MHC, Hoescht, or Edu were taken with a 10× objective with at least 25 stacks at 10 uM spacing. Magnified images of nuclei in the myocardium of PHTs were taken with the 40xW objective with 30 slices at 1 uM spacing.

### 5.2 Image Processing and Analysis

Images were stitched and analyzed using Fiji(ImageJ)(Preibisch, Saalfeld, and Tomancak, 2009). The cell counting is automated with the help of a state-of-the-art deep learning technique. We train an artificial intelligence (AI) system to predict high scores near the centroid of each chicken nucleus in a 2D image (Figure 4). In order to avoid the curse of dimensionality, we have converted the task of nuclei counting in 3D image stacks into a nuclei detection problem in 2D projection of image slices. We first compute a maximum projection of slices from 3D confocal image stack. The nuclei locations in the 2D projected image are annotated by the biologist. Corresponding to the max-projected image, a target matrix is prepared by placing small gaussian peaks at the annotated center locations of the nuclei in a zero filled matrix(Hughes *et al*., 2018; Xie, Noble and Zisserman, 2018). The size of this matrix is equal to the size of the max-projected image. The problem of nuclei detection then reduces to matrix regression in a two-dimensional space. A U-net style autoencoder(Ronneberger, Fischer and Brox, 2015) is trained to regress the gaussian scores in response to the 2D input image in order to detect the nuclei centers. Subsequently, a 2D peak detection algorithm (Van Der Walt *et al*., 2014) when applied on the output scoremap produces the estimated locations of the nuclei. This method works because the nuclei distribution in 3D shifts slice wise so that their projection does not lead to major occlusion problems. Additional details can be found in the supplementary material. Our representative implementation of the cell counting methodology is available at: https://github.com/CCCofficial/cell-counting-chicken-embryo-myocardial

### 6.1 Statistical methods

Statistical significance was determined by the one-tailed *t*-test. All statistical analysis was performed with Prism 7 (Graphpad) and *P* values are reported (n.s. = p > 0.05, * = p ≤ 0.05). All error bars represent SEM.

## Notes

https://github.com/CCCofficial/cell-counting-chicken-embryo-myocardial

### Competing Interest Statement

The authors have declared no competing interest.

