## Supplemental Figures for "Endoderm Nitric Oxide Signals to Regulate Nascent Development of Cardiac Progenitors in Chicken Embryos"

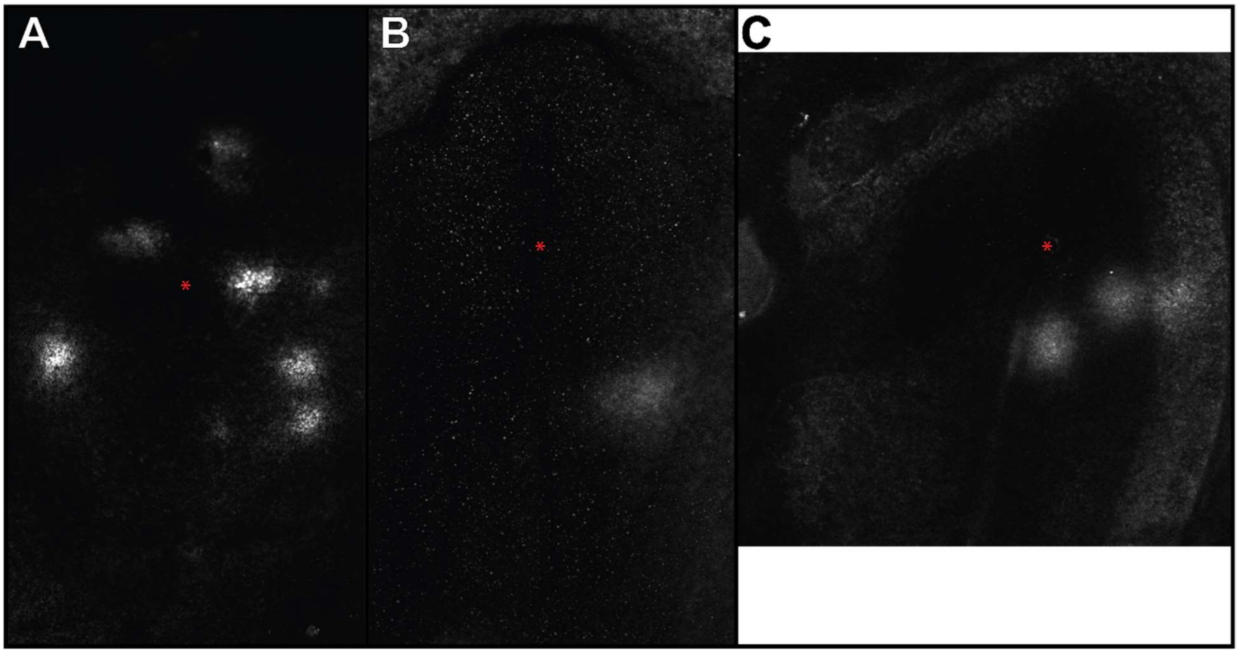


**Sup. Figure1. Live DAF2-DA labeled HH4 Embryos.** Live confocal fluorescence images of three age matched HH4 staged embryos as shown in composite image Figure 2C. Ventral sided DAF2-DA labeled embryos acquired with (A, B) a 10X objective and (C) a 20X objective lens. NO hotspots lateral of Hensen's node, indicated by "*", are within regions where cardiac precursors are migrating laterally to for the cardiac mesoderm suggesting that NO may be involved in these cellular events.

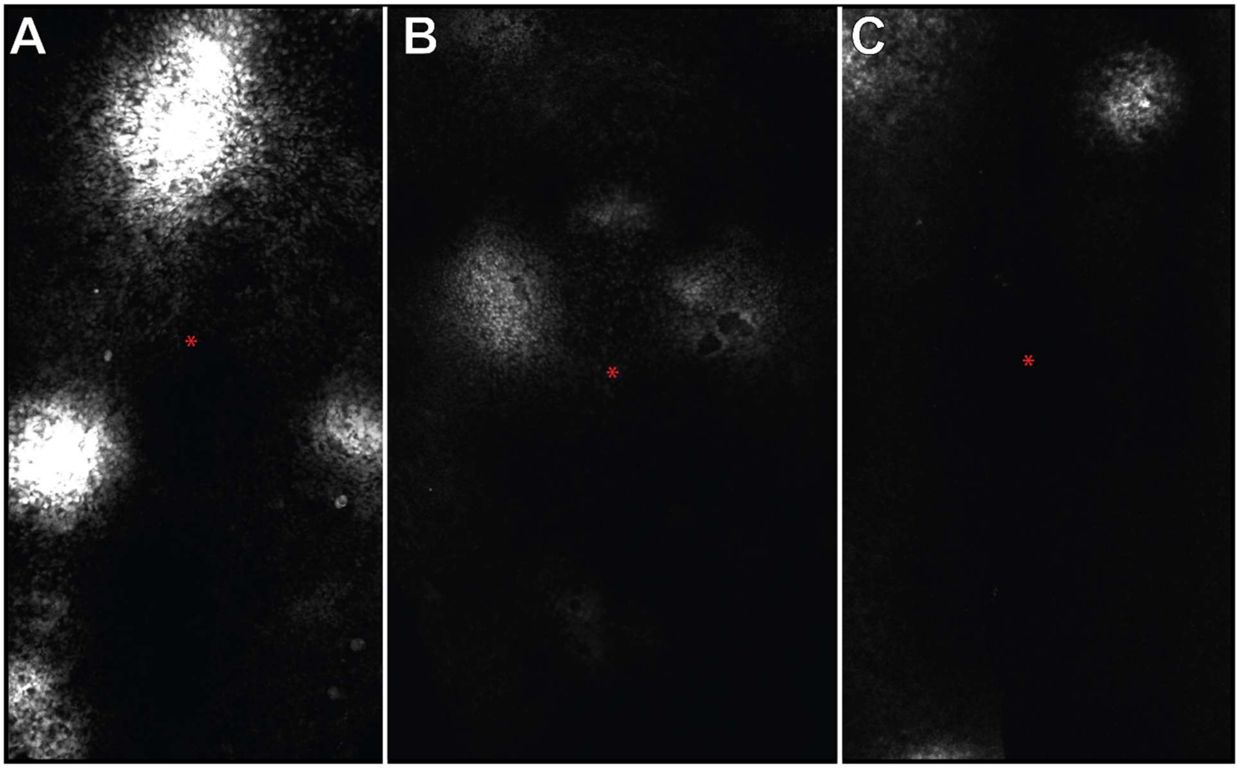


**Sup. Figure2. Live DAF-2DA labeled HH5-6 Embryos.** Live confocal fluorescence images of three age matched HH5-6 staged embryos as shown in composite image Figure 1C. (A-C) Ventral sided DAF-2DA labeled embryos acquired with a 10X objective and stitched vertically using ImageJ. The NO hotspots persists at this stage of development and continues to be lateral of Hensen's node (*). These hotspots are in regions of the embryo associated with the migration of precursors that are forming the arc of the cardiac crescent in HH5-6 embryos.

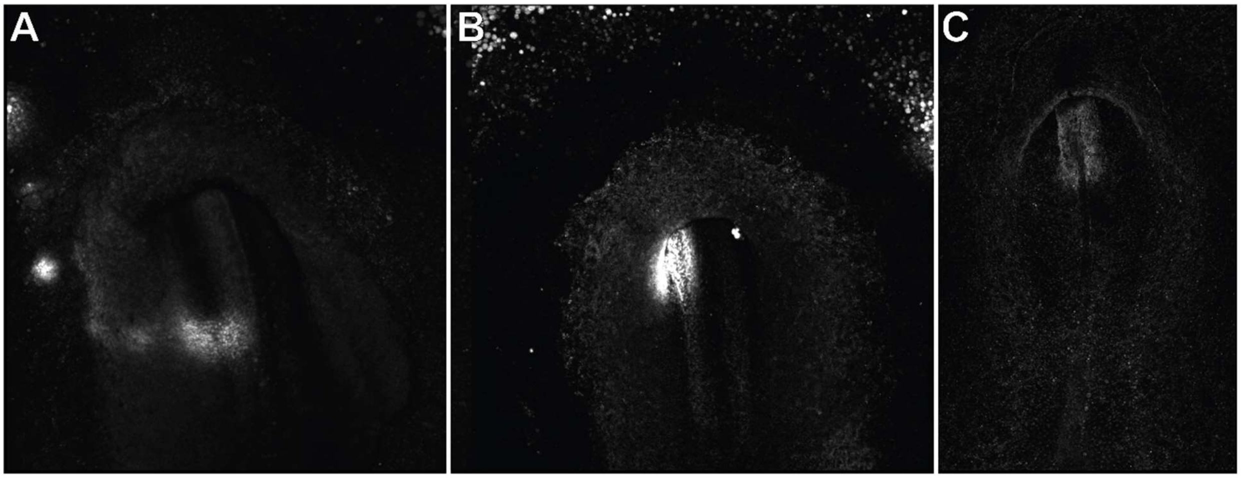


**Sup. Figure3. Live DAF labeled HH7-8 Embryos.** Live confocal fluorescence images of three age matched HH7-8 staged embryos as shown in composite image Fig 2G. (A-C) Ventral sided DAF-2DA labeled embryos acquired with a 10X objective. (A, B) NO hotspots in regions concentrated around the secondary heart field where migrating precursors begin to fuse and form into the endocardial tube. (C) Exhibits a NO concentrated over the region of the endoderm where the endocardial tube is forming.

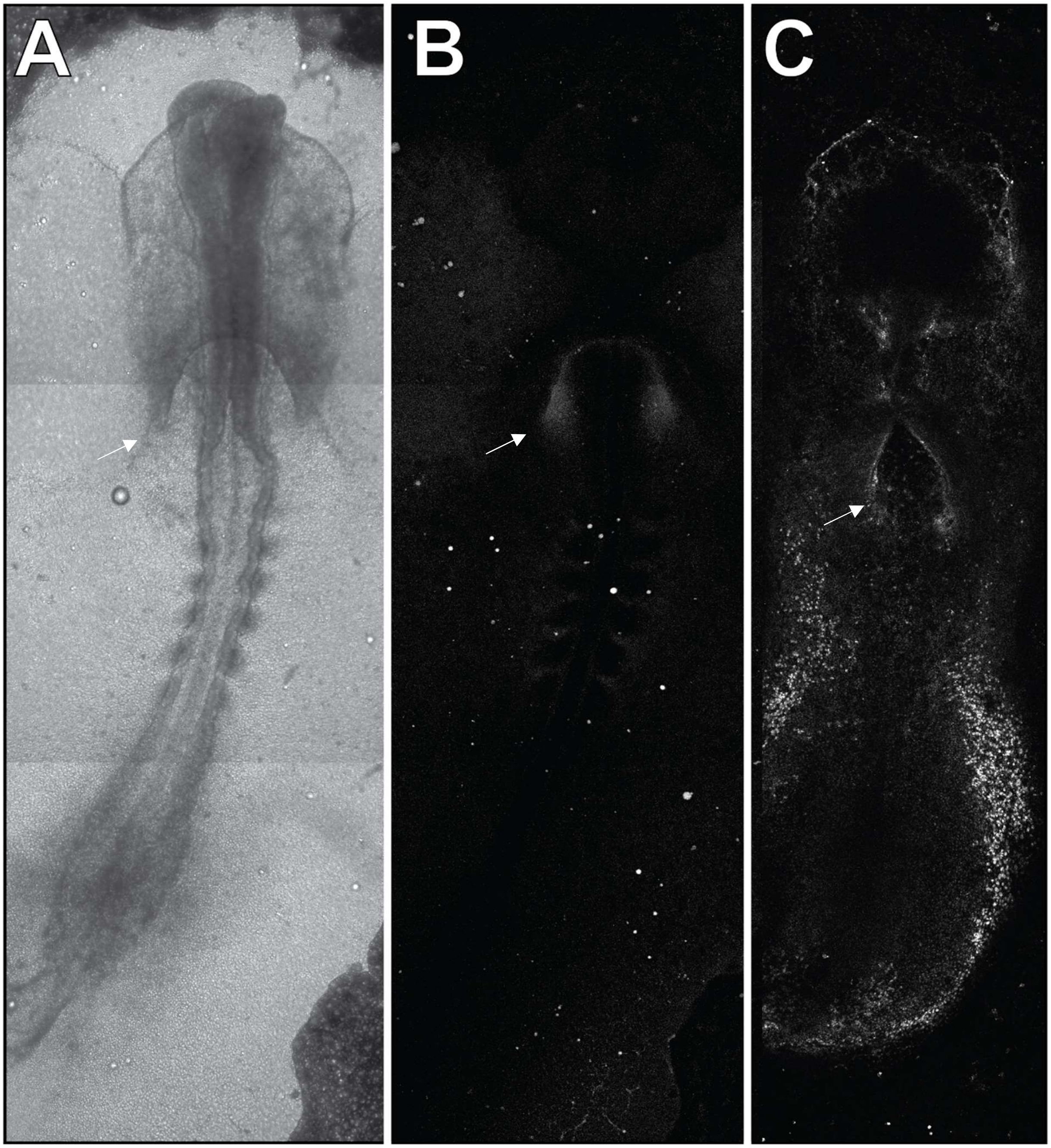


**Sup. Figure 4. Live DAF2-DA labeled HH8-9 Embryos**. Live confocal fluorescence images of age matched HH8-9staged embryos. Ventral sided images of (A) transmitted light and (B, C) DAF2-DA labeled whole embryos. Images acquired with 10X objective and stitched vertically using ImageJ. (B, C) NO hotpots shown in the secondary heart field regions where progenitor cells are adding to the forming primitive heart tube (arrows).

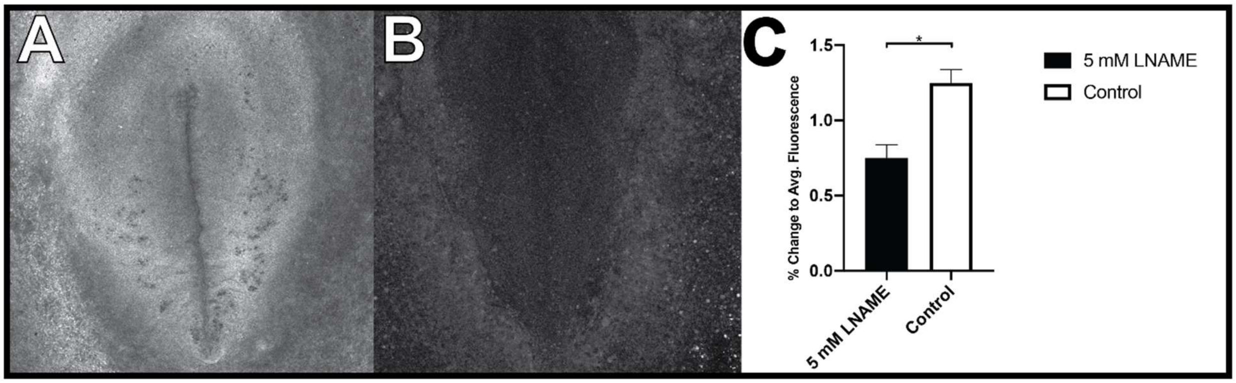


**Sup. Figure 5. In ovo endoderm L-NAME treatment inhibits NO formation.** Confocal DAF2-DA fluorescence image of (A) Control and (B) 5 mM L-NAME treated embryo. (C) 1 hour endoderm treatment significantly attenuated NO signaling in HH4-5 staged embryos. ( -39.9± 0.08%, n=6,p=0.01). 1-tailed student's T-test. (*p< 0.002). Demonstrates the ability to target endoderm NO formation in the developing embryo via sub-germinal cavity injection.

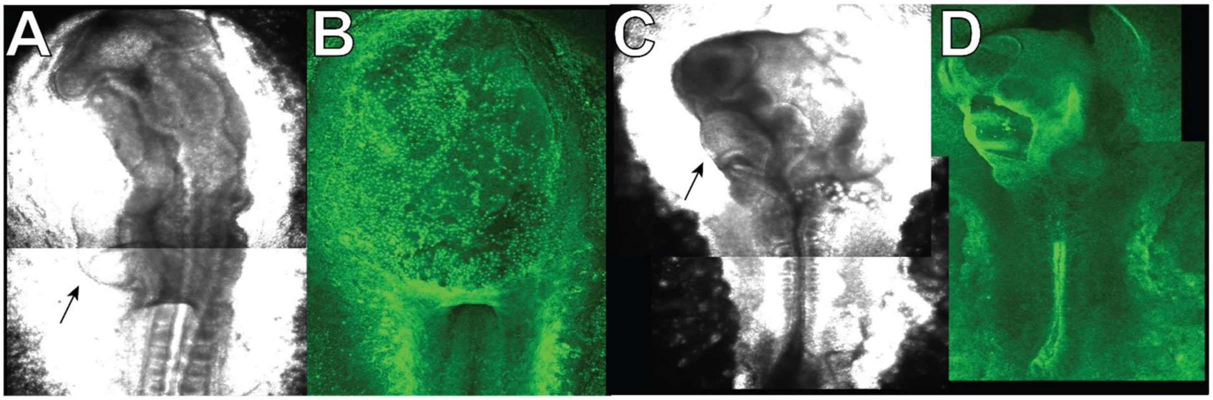


**Sup. Figure 6. Strong NOS inhibition induces severe morphological defects** Transmitted light images (A,C) of HH11-12 staged embryos treated with (A, B) Vehicle and or (C,D) 25mM LNAME. Severe morphological defect in the PHT (Black arrow) seen in (C) LNAME treated embryo compared to (A) control. Demonstrates a requirement of NO signaling for normal development of the embryo.
