## Supplemental Text for "Endoderm Nitric Oxide Signals to Regulate Nascent Development of Cardiac Progenitors in Chicken Embryos"

**Supplemental Materials**

**S1. Introduction**

A computational approach to cell counting is studied to investigate the cell proliferation in the confocal immunofluorescence images of cardiac tissues of chicken embryos. The image dataset consists of a total of 15 three-dimensional image stacks. Each stack is made of 30 slices where each slice is a single channel with dimension 512×512 pixels. The nuclei are arranged in the stack in such a fashion that if one takes the maximum projection of the slices, almost all of the nuclei remain fairly visible in the projected view, which means that the nuclei can be identified, and counted, in the 2D view.

**S2. Problem Statement and Computational Tasks**

We propose the nuclei counting problem as counting-by-detection. That means, – the nuclei are detected first, and then counted. There are of course alternate approaches. For example, one can sum the predicted density function in a given region in order to estimate the number of nuclei (Xie *et al.,* 2016). Another approach could be designing an end-to-end deep learning network that directly delivers the count as output (Cohen *et al.,* 2017). However, we have followed a relatively simple and straightforward method of counting-by-detection in this paper.

To train a machine learning system one requires high quality annotation. To that end, an expert biologist in our group has annotated the centers of the chicken nuclei using the widely used software ImageJ/Fiji. The annotation is done in each 3D stack using the multi-point tool of the Fiji software. This results in a set of 3D center coordinates each corresponding to a particular nucleus in the image stack.

Any machine learning algorithm that works directly with 3D data requires a relatively larger system configuration. Besides bigger GPUs and larger memories, the time required for processing the higher dimensional data is also high. Our experience suggests that ensuring a stable gradient during the backpropagation is also a little tricky with three-dimensional data. We advocate a simple, efficient and effective system, and to facilitate that we have decided to reduce the data dimensionality from 3D to 2D. In principle, we have taken the maximum projection of intensities in the image stack along Z-direction. This process preserves the height and width of the image slices in the resulting image, but the number of Z-slices is reduced to one. Accordingly, we ignore the Z-coordinate in the annotation and consider only the x and y coordinates to refer to the nuclei centers.

Converting 3D to 2D may result in information loss along the Z-direction, particularly nuclei occlusion. In our data, the nuclei distribution in each slice is slightly shifted in xy plane relative to its immediately preceding or immediately succeeding slice. When projected on a single plane we observe limited occlusion. The instances of partial overlap do not significantly reduce the extracted information when projected. Fig. 1(a) shows five uniformly sampled slices and one can see different spatial information in the xy-plane that gets highlighted in different slices. The resultant max-projected slice on the right summarily captures the entire nuclei distribution in the full 3D stack.

**S2.1. Detection as High Dimensional Matrix Regression**

At this point, our problem statement reduces to identifying nuclei in 2D (max-projected) image slices. We also have the 2D coordinates of these nuclei that serve as ground truth. However, the number of image slices (only 15) is not large enough for today’s state of the art machine learning. We solve this problem by splitting each max-projected slice in four equal sized image tiles. This leads to a sufficient number of images to carry out our machine learning training process (see Fig. 1(b)).

We have formulated the machine learning problem as a matrix regression task. Corresponding to each image input we create a zero filled matrix of the same size. The nuclei’s center coordinates in the zero matrix are filled with small Gaussian weights that model the location likelihood of the nuclei centers. This matrix serves as the target matrix in our matrix regression task. In response to an image tile as input the task of our deep learning system is to regress the target matrix (Fig. 1(c)). The predicted weights (i.e, estimated Gaussians) in the output matrix would indicate the location likelihood of the nuclei centers.

**S3. Methodology**

This high dimensional matrix regression task illustrated in Fig. 1(c) is quite general. In principle, any autoencoder-decoder architecture can be used for this purpose. In this work we have proposed to use the U-Net style autoencoder-decoder to perform the matrix regression. U-Net is originally proposed to perform object segmentation (Ronneberger *et al.*, 2015). Following its excellent performance and remarkable predictive power in high dimension, subsequent studies have extended the U-Net to other machine learning tasks like density prediction (Xie *et al.,* 2016) and detection (Hughes *et al.,* 2018). We take inspiration from (Hughes *et al.*, 2018) and apply this technique to address the current problem statement. Converting the problem statement from 3D to 2D makes it readily amenable to this kind of matrix regression.

**S3.1. Neural Architecture and Training**

The U-Net architecture is illustrated in Figure 2(a). The usual cross-entropy loss function of U-Net (Ronneberger *et al.*, 2015) is changed to MSE-loss (Xie *et al.,* 2016; Hughes *et al.*, 2018). The back-propagated error gradients gradually adjust the weights of the network helping it learn to produce output matrices increasingly similar to the target matrices. Each layer in the convolutional neural net in Fig. 2(a) comprises certain fundamental units like *C*(*m*, *n*) that indicates m input channels producing n output channels. The default size of the kernel used in the convolution is 3.

Though reducing the data dimension and increasing the number of images helps the training task, additional data augmentation certainly improves the training performance. Recent reports suggest that the training with deep learning architectures benefits from artificially augmented datasets (Krizhevsky *et al.,* 2012). Such augmentation adds diversity to the existing set of images in the form of various transformations. As a consequence, the training algorithm gets to see more variations that often result in better performance. We have followed the image transformations indicated below, and applied them to both input and the target matrices:

i. Horizontal and vertical flipping
ii. Horizontal/vertical shifting by a maximum of 30% width/height
iii. Zooming in/out randomly up to 30%
iv. Rotation randomly by up to 30 degrees

Each of the geometric transformations above happens in a random fashion and is implemented as a part of the data loading module in the deep learning library (Paszke *et al.*, 2018). We select a transformation randomly with a probability of 1/2. Also, we select the parameters of the transformation from the corresponding uniform distribution defined for each parameter (Xie *et al.,* 2016).

The training and validation losses are monitored as a function of iterations. We observe that the training converges in about 100 iterations as shown in Fig. 2(b).

In order to reduce various experimental biases that might have influenced data acquisition and preparation, we have resorted to a K-fold cross validation technique. An alternative approach to split the dataset into a one-time train and test partitions and reporting results only on that particular partition may favor the outcome in a wrong way. With the K-fold cross validation technique we have the opportunity to do the evaluation multiple times while reporting results on the aggregated output.

In particular, we have used a 5-fold cross validation technique. Each time we leave out 3 image stacks for testing while training and validating on the rest. This allows us to rotate the testing set, eventually covering the entire dataset. At the end of the experiment, we combine the test results from all folds obtaining the predictions on all of the 15 image stacks. We use the test results for extensive evaluation of the methodology (details given in the Results section).

**S3.2. Inference**

Since we use image tiles as input to the autoencoder the inference step requires some extra steps. Each test image is split into four equal sized image tiles as shown in Fig. 3. The resulting tiles are processed by the trained autoencoder. The output scoremaps are collected and placed in the appropriate grid position to build the prediction scoremap for the full image.

A peak detection algorithm (Walt *et al.*, 2014) operates on the scoremap identifying local peaks. The location coordinates of the local peaks serve as estimated nuclei centers. The algorithm (Walt *et al.*, 2014) has one important parameter that we have used to control our output, namely, minimum distance, represented by *d*. This distance actually specifies a window defined as follows:

*W* = 2 x *d* + 1	(1)

Within this window the secondary peaks, if any, would be made zero permanently. This is an important and essential post-processing step called non-maximum suppression and is widely used in the detection literature (Hosang *et al.,* 2017).

**S4. Results and Discussion**

The predicted scoremaps and the subsequent peak detection deliver the final coordinates of the nuclei centers. The estimated nuclei centers are compared with the ground truth to ascertain the effectiveness of the methodology we have followed in this work. Specifically, the problem of comparing the predicted coordinates with the ground truth ones can be addressed by point set matching. Two points sets coming from two sources can be matched well by Hungarian algorithm. We have used such techniques to associate the points sets in order to find out correct and incorrect predictions.

Essentially, we have computed the three-performance metrics. But before we discuss the performance metrics few important terminologies are worth mentioning in the context of our work. When the predicted coordinates match with any of the ground truth we call this prediction as true positive. The prediction that fails to find a match with the ground truth is termed as false positive. Lastly, the ground truth that goes unmatched with any of the predictions is referred to as false negative. Based on these notions we define the performance metrics as follows:

precision = true positives / (true positives + false positives) 	(2)

recall = true positives / (true positives + false negatives)   	(3)

The precision refers to the percentage of predictions that is correctly classified. In contrast, the recall represents the percentage of ground truth that is correctly classified. The last metric that combines these two and thereby proving a single metric for evaluation is F-score which is actually the harmonic mean of the precision (2) and recall (3).

Figure 4 shows the performance evaluation. The peak finding algorithm requires an input distance *d* (1) for applying non-maximal suppression. This parameter d is important for ensuring the desirable characteristics of the proposed methodology. To facilitate setting *d* to the right value we have followed a data driven approach. In principle, we have varied *d* over a range of 1 to 8, and for each value of *d* we compute the matches, misses, and spurious detections leading to a set of precision, recall and F-score. This characteristic is illustrated in the plot of Fig 4(a). Note that with the increase in value of *d* the precision improves but the recall subsides. This tradeoff has a “sweet spot” at around *d* = 3 when both the values are the same. We set *d*  to this value as our operating point to report the final performance measures as shown in Fig. 4(b).

A visual comparison between the ground truth and the predicted nuclei centers is produced in the Fig. 5.  The annotated nuclei centers are marked in red in the first column and the predicted centers in blue in the last one. The middle columns show the scoremaps, the likelihood distribution for the nuclei centers as delivered by the neural autoencoder. It is clear to see that the trained autoencoder has learned the generalization pretty well. Even in case of crowded nuclei clusters the model has predicted the likelihood score to an excellent degree, meaning that the local likelihood peaks capture the shape of the nuclei clusters pretty well. This shows the sensitive nature of our algorithm. The model performs pretty well in terms of specificity. Empty spaces in the image are correctly identified as empty in the prediction scoremap, and no spurious predictions are seen in these regions. Also, the neural network is seen to make conservative predictions because the nuclei centers when blurry do not excite the neural net to fire on them.

**S5. Conclusion**

In this work the objective is to count the nuclei centers in 3D confocal microscopy. Given the small number of input images we have followed an aggressive dimensionality reduction and data augmentation strategy to avoid overfitting. The counting task is approached from a counting-by-detection perspective and the detection problem is formulated in terms of high dimensional matrix regression. Such regression task is carried out with a U-Net style autoencoder that predicts scoremaps in response to an input image. The scoremaps are processed by a peak finding algorithm to estimate locations of nuclei centers.

The evaluation of this proposed technique shows good generalization capability. We have supported our findings with a series of experimental results. We believe this methodology will introduce a new way of analyzing and quantifying 3D data, particularly in the context of cellular object detection and counting. Our representative implementation of the cell counting methodology is available at: <https://github.com/CCCofficial/cell-counting-chicken-embryo-myocardial>

**Acknowledgements**

We thank Lisa Galli for providing pivotal guidance in experimental procedures, as well as generously providing antibodies and laboratory tools used in this study. We also thank Dr. Takashi Mikawa for constructive discussions and his expertise in the field of embryonic cardiology. We thank Annette Chan and the Cell Molecular Imaging Center at SFSU for technical support with microscopy. Y.-H. M. C. and W. L. C. acknowledge funding from NIH award 1SC2GM118267 and the NSF STC Center for Cellular Construction award DBI-1548297. S.B. and S. K. B also thankfully acknowledge the NSF STC Center for Cellular Construction award DBI-1548297.

**Figures**

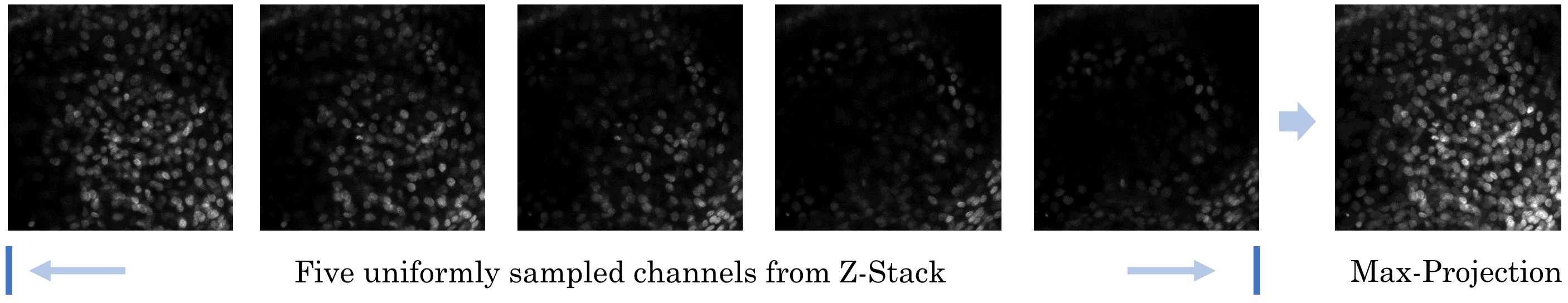

(a)

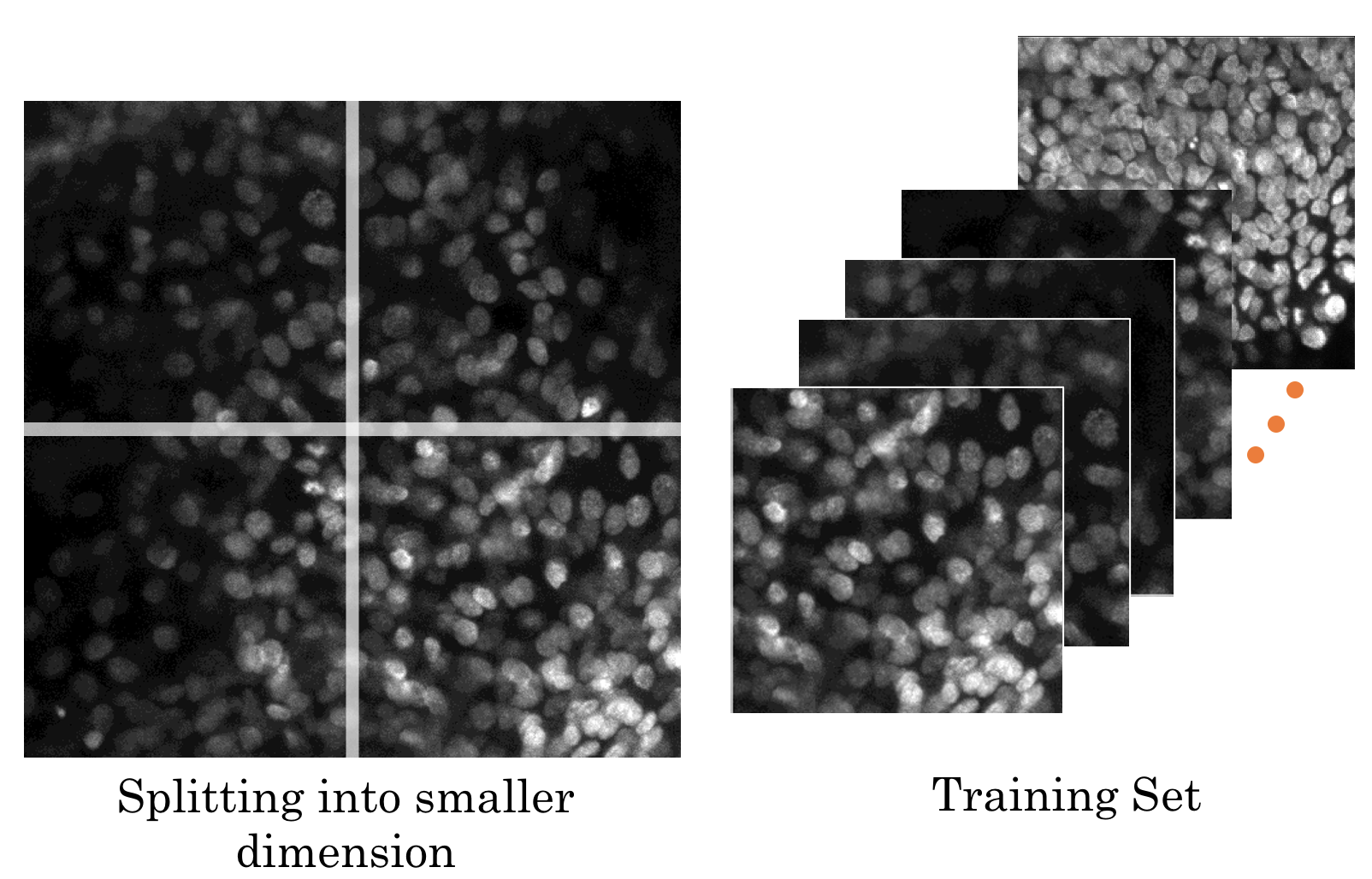

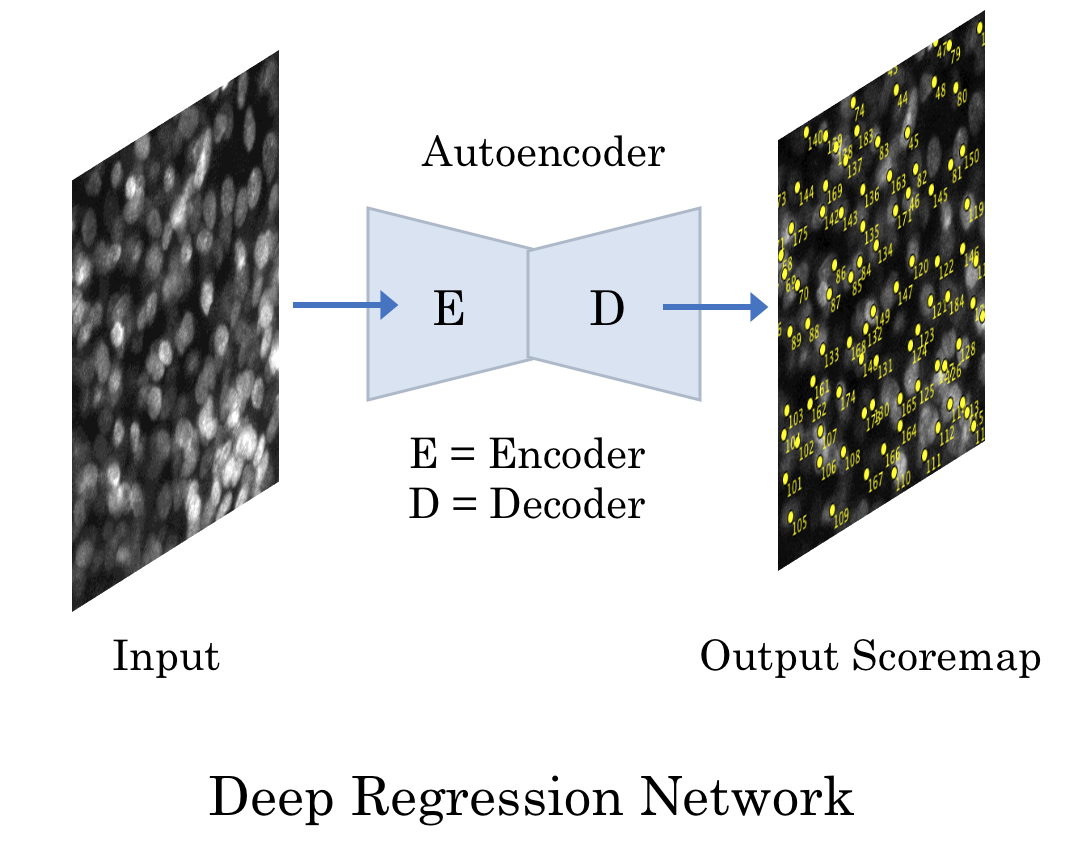

(b)	(c)

**Figure 1** (a) One of the 3D confocal image stacks is shown with five uniformly sampled channels (or slices) in the z-stack. Each stack has 30 slices and we apply a max-projection to convert the computer vision problem into 2D. (b) We further reduce the dimensionality of the problem by splitting the 512 x 512 max-projection into four equal image tiles as illustrated. We train the machine learning algorithms with this set of images. (c) Training The target matrix comprises tiny Gaussian blobs in the place of nuclei centers. A U-Net style autoencoder learns to regress these Gaussian weights as a likelihood of nuclei centers. The deep regression network directly operates in the image space, meaning the network takes the image tiles as input from (b) and produces an output score matrix that has the same size as that of the input image tiles in (b). Consequently, it becomes straightforward to draw direct correspondence between predicted gaussian weights and the cell centers in the input image tiles, without having to do any coordinate transformation.

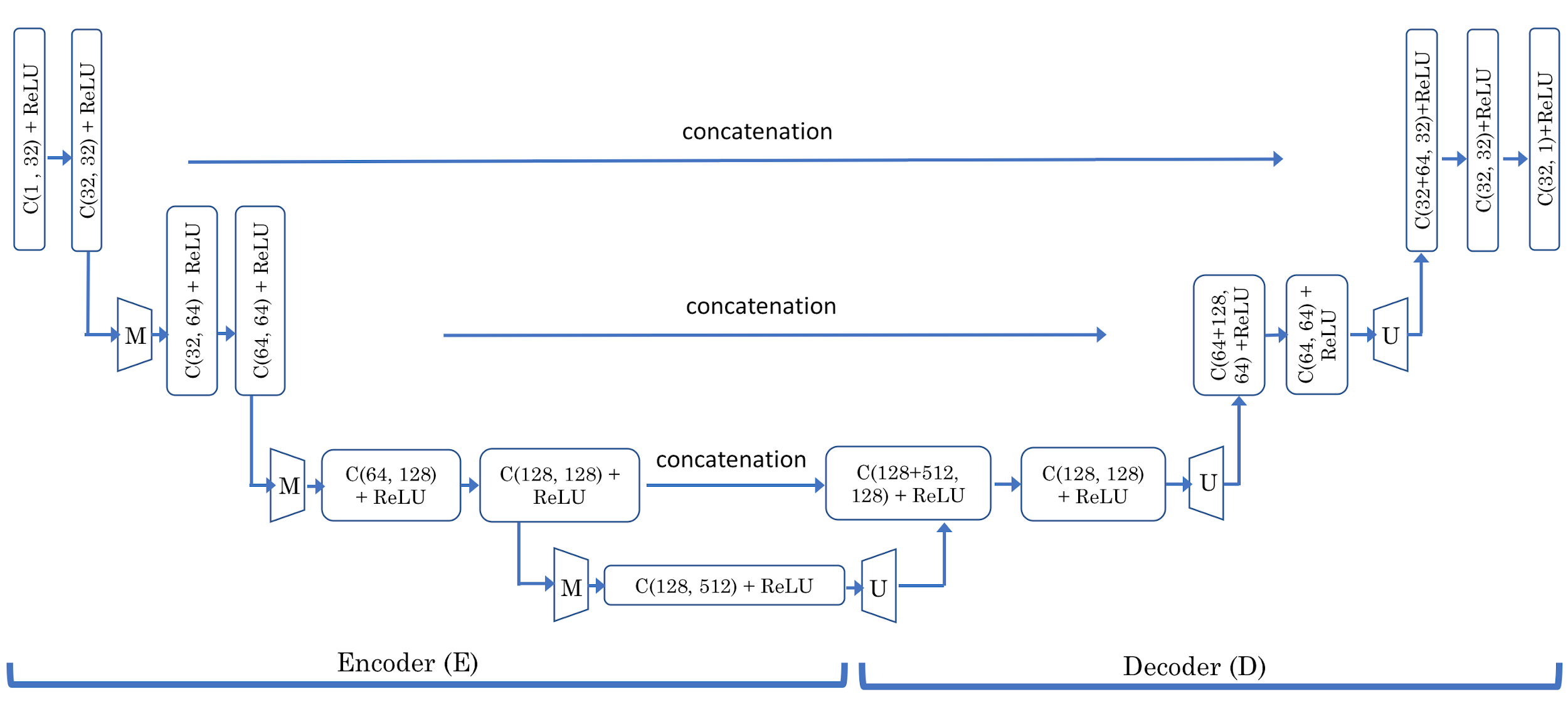

(a)

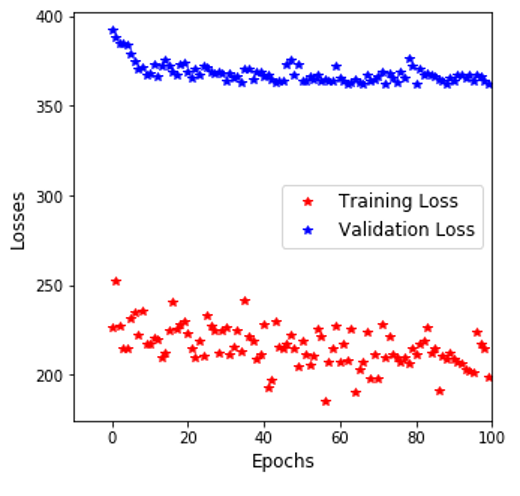

(b)

**Figure 2** (a) The neural net architecture is shown with its two primary components, namely, the encoder and the decoder part. The encoder part gradually reduces the size of the image as a result of the maxpool operation, whereas the decoder module does the opposite to bring the resultant output in its original size. The encoder (E) and decoder (D) parts of the neural architecture are shown with all their constituent layers. Note, C(m, n) actually denotes an entity comprising three individual components: a convolutional layer having m input channels and n output channels with a kernel size of 3, a rectified linear unit (ReLU), and a batch-normalization unit. The max-pooling (denoted by M) has a default size of 2 × 2, and the upscale factor (denoted by U) has a default value of 2. The loss function imposed on the output layer is MSE (mean squared error) loss. (b) Training and validation losses incurred by our network architecture are shown during the training process as a function of the iteration number.

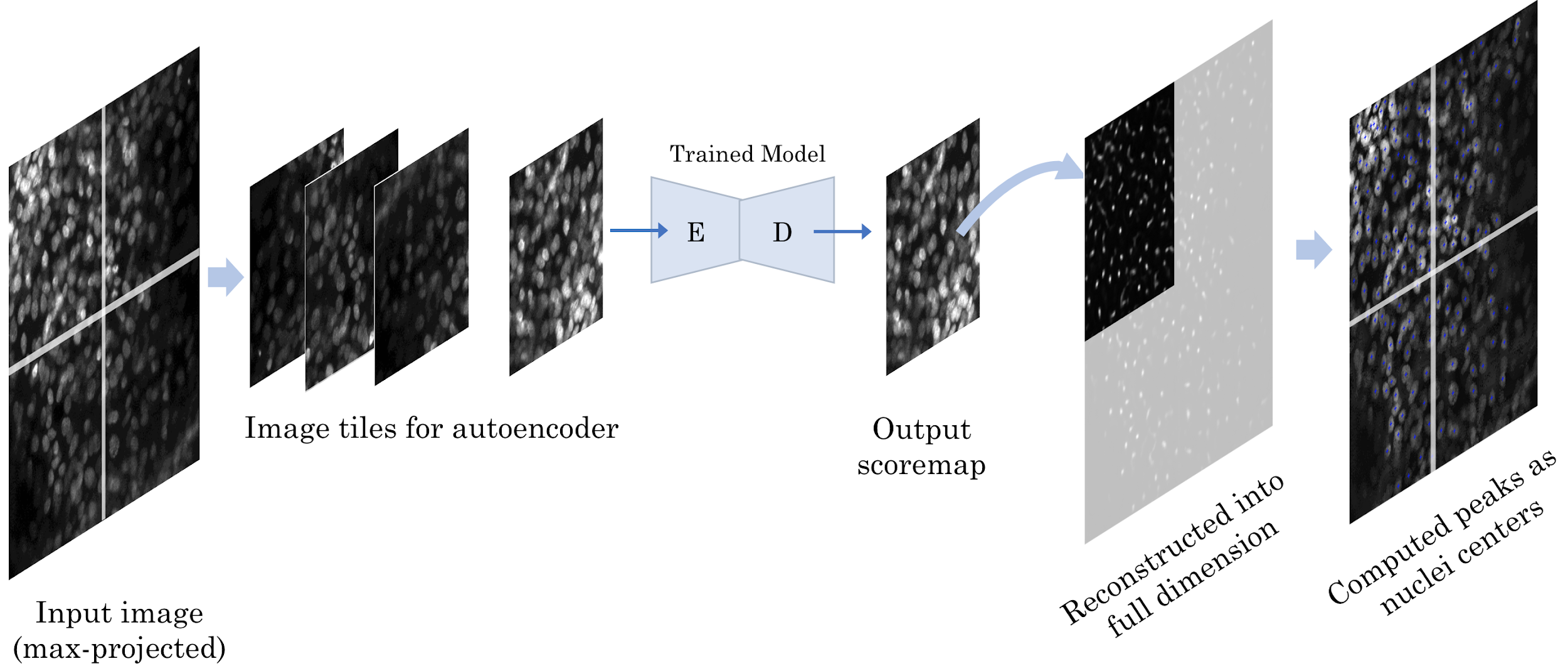

**Figure 3** In the Inference stage the objective is to predict the nuclei centers using the trained U-Net autoencoder. The max-projected image derived from the 3D image stack is split into four tiles as shown on the extreme left. The four tiles are independently processed by the trained model to yield detection scoremaps. The output tiles are placed in the respective positions in the original image grid reconstructing the likelihood map corresponding to the original full-scale image. A local peak detection algorithm processes the likelihood scoremap computing the nuclei centers as the final output.

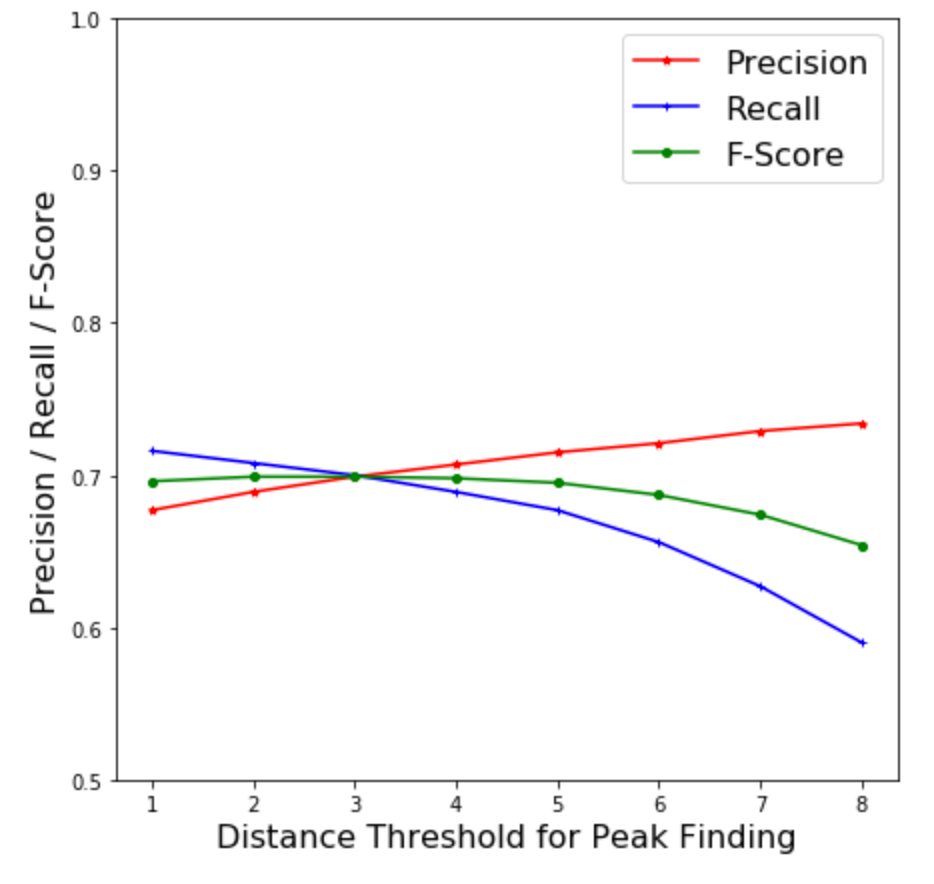

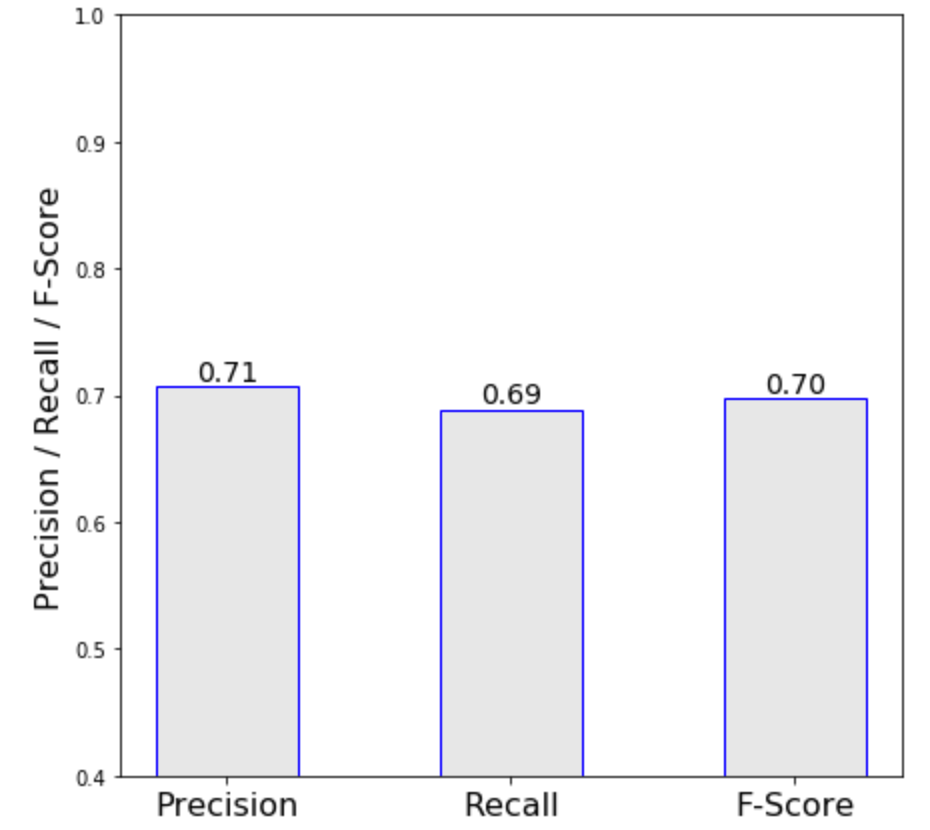

(a)	(b)

**Figure 4** Performance evaluation (a) The neural net delivers the prediction scoremap which is actually an estimation of gaussian weights in the target matrix. The predicted scoremap is processed by a peak-finding-algorithm; the algorithm requires a distance input as a parameter. Within this distance any secondary peak besides a primary one will be removed permanently. Varying this distance varies the prediction. The varying precision, recall, and F-Scores are exhibited that require Precision as a function of varying distance threshold. Note, with the gradual increase in this distance parameter precision increases but the recall falls. The sweet point where both are approximately the same marks our operating zone. We decide the final value of this threshold as the point where the precision and recall curves cross each other. (b) At the computed threshold we show the final precision, recall and F-Score in this plot.

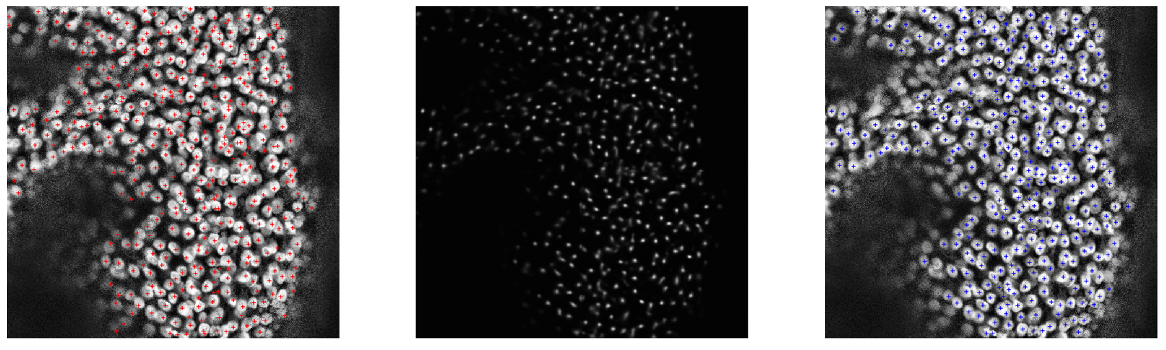

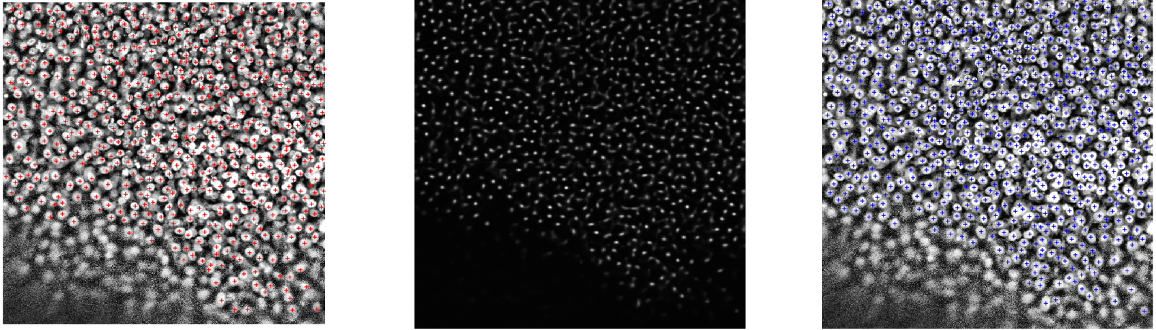

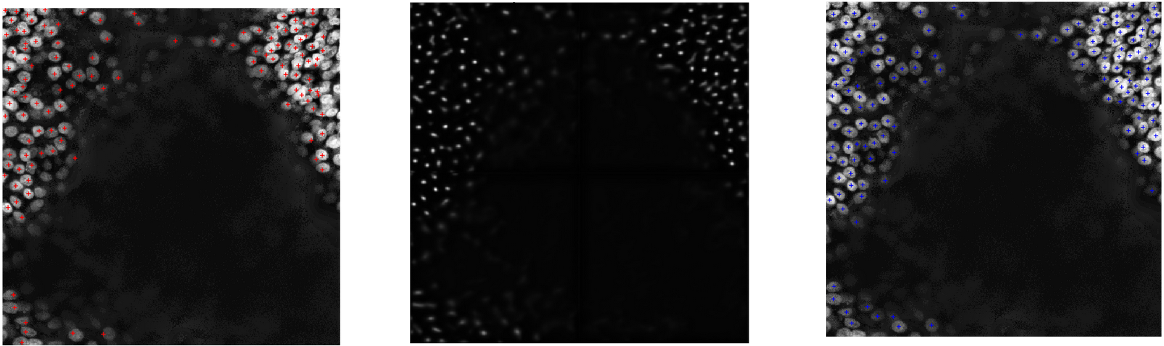

Ground truth overlaid	Predicted scoremap	Detected peaks overlaid

**Figure 5** The left column shows ground truth overlaid (in red) on the full-sized max-projection. The middle column shows the predicted scoremaps with regressed Gaussian weights. Note how nicely the peaks in these scoremaps correspond to the nuclei centers in the left column. A peak detection algorithm identifies the local peaks, and their coordinates represent the estimated nuclei centers. The third column shows predicted nuclei centers (in blue) overlaid on top of the max-projected image.
